# Landscape of the Dark Transcriptome Revealed through Re-mining Massive RNA-Seq Data

**DOI:** 10.1101/671263

**Authors:** Jing Li, Urminder Singh, Zebulun Arendsee, Eve Syrkin Wurtele

**Affiliations:** Genetics and Genomics Graduate Program, Iowa State University, Ames, 541004, USA; Bioinformatics and Computational Biology Program, Iowa State University, Ames, 50014, USA; Department of Genetics, Development, and Cell Biology, Iowa State University, Ames, 50014, USA; Center for Metabolic Biology, Iowa State University, Ames, 50014, USA

**Author notes:** Corresponding author. (Wurtele ES).

**Keywords:** orphan gene, *de novo*, RNA-Seq, Ribo-Seq, gene function, cluster analysis

## Abstract

The “dark transcriptome” can be considered the multitude of sequences that are transcribed but not annotated as genes. We evaluated expression of 6,692 annotated genes and 29,354 unannotated ORFs in the *Saccharomyces cerevisiae* genome across diverse environmental, genetic and developmental conditions (3,457 RNA-Seq samples). Over 48% of the transcribed ORFs have translation evidence. Phylostratigraphic analysis infers most of these transcribed ORFs would encode species-specific proteins (“orphan-ORFs”); hundreds have mean expression comparable to annotated genes. These data reveal unannotated ORFs most likely to be protein-coding genes. We partitioned a co-expression matrix by Markov Chain Clustering; the resultant clusters contain 2,468 orphan-ORFs. We provide the aggregated RNA-Seq yeast data with extensive metadata as a project in MetaOmGraph, a tool designed for interactive analysis and visualization. This approach enables reuse of public RNA-Seq data for exploratory discovery, providing a rich context for experimentalists to make novel, experimentally-testable hypotheses about candidate genes.

## Introduction

Pervasive transcription of unannotated genome sequence in eukaryotic species is evidenced in multiple RNA-Seq studies. [1–5]. Indeed, transcription and translation has been described for non-genic regions of genomes in diverse species [6–15]. Many studies have dismissed this expression as transcriptional “noise” [4, 16–18]. However, several functional genes have been identified from the so-called “noise” [19, 20]. This mass of unannotated transcripts, often ignored and little understood, we refer to as the “dark transcriptome” (Figure 1.A).

**Figure 1.**
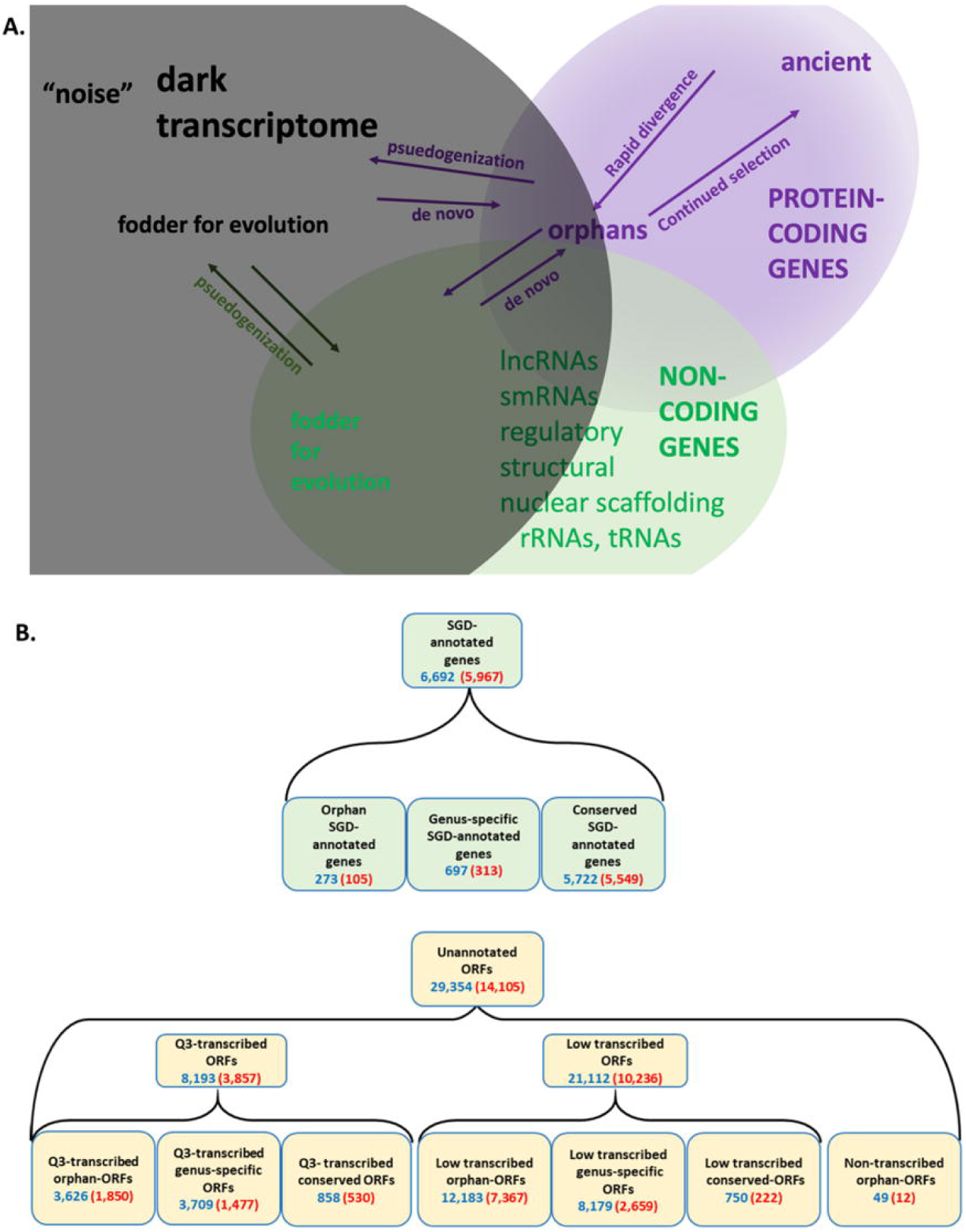
Quantification of SGD annotated genes and dark transcriptome. **A.** Definition of Dark transcriptome. Pervasive transcription of unannotated sequences has been found in many species. Some of these might be protein coding genes that have escaped annotation. Most of these unannotated coding genes are orphan (species-specific) genes, which have no homolog to other species, and are hard to predict using current gene prediction tools. These orphan genes could emerge by rapid divergence from ancient genes or could evolve *de novo*. Other transcribed but unannotated sequences might be non-coding genes. Although many studies have explored the function and classification of the non-coding transcripts, many transcribed sequences are still unclassified. **B.** Classification and numbers of expressed transcripts for SGD-annotated genes (green boxes) and ORFS (yellow boxes). Orphan-ORFs, unique to *Saccharomyces cerevisiae* (phylostrata (PS)=15); genus-specific-ORFs, unique to *Saccharomyces* spp. (PS=10-14); conserved-ORFs, homologs in older species (PS=1-9). Q3-transcribed, ORFs with mean transcription across the 3,457 samples ranking in the upper (Q3) quantile of the unannotated transcripts. Low-transcribed ORFs, ORFs with mean transcription across the 3,457 samples ranking in the lower 75% of the unannotated transcripts. Non-transcribed orphan-ORFs (Figure S8). Red font, number of genes/ORFs with translation evidence according to Ribo-Seq analysis. (For full PS designations and transcription expression, see supplementary file, *S. cerevisiae_RNA-seq_3457_27.mog*; for translation per transcript, see supplementary file, *Ribo-Seq_rawcounts.csv.*)

Each organism contains species-specific genes (denoted here as “orphan genes”). The challenge of distinguishing orphan genes in genomes and predicting their functions is immense, resulting in an under-appreciation of their importance. The emergence of novel protein coding genes specific to a single species (orphans) is a vital mechanism that allows organisms to survive a changing environment [21, 7, 22–25]. Over generations, those orphan genes that continue to provide a survival advantage will be maintained. Orphan genes can be identified from within a list of genes by phylostratigraphy, the classification of each gene according to its inferred age of emergence [21, 22]. Two general mechanisms enable orphan gene emergence:1) *de novo* evolution and 2) divergence.

Orphan genes can evolve *de novo* from non-coding sequence in regions of the genome lacking genes entirely or as new reading frames within existing genes [11, 22, 26, 27]. Indeed, transcriptional and translational “noise” has been suggested as a mechanism that facilitates novel gene emergence [28–33]. This hypothesis is borne out by *in vitro* and *in vivo* synthetic biology research demonstrating that novel peptides are often able to bind small molecules (e.g., ATP, and metals) [34] and induce beneficial phenotypes when expressed [34, 35]. If information on the expression of the dark transcriptome was more easily accessible, the potential roles of expressed transcripts could be better considered.

Orphan genes can also evolve from existing proteins by divergence of protein coding sequences (CDSs) beyond recognition [22, 28, 29, 31, 33, 36–39]. We estimate from the phylostratigraphic data on yeast genes that this process would require ultra-rapid sequence divergence relative to that of the average protein. Evolution of orphan genes from existing protein-coding genes has been estimated to account for about 18% (human), 25% (Drosophila), and 45% (yeast) of annotated taxonomically-restricted genes [39]. (This estimate considers only the ∼50% of yeast genes that can be compared across species, i.e., those that are located within syntenic intervals of related genomes [40].)

A systematic analysis of current computational methods for genome annotation indicates many orphan genes may be missed in annotation projects [41]. This is because genes are often identified from sequenced genomes by combining evidence based on homology with other species [42, 43] with *ab initio* machine-learning predictions by detecting canonical sequence motifs (e.g., splice junctions) [44, 45]. However, homology and *ab initio* approaches can be problematic in predicting orphan genes. First, orphan genes cannot be identified by homology to genes of other species, since they have none. Secondly, to the extent that an orphan has not yet evolved canonical motifs, *ab initio* prediction may be ineffective. For example, compared to the gold-standard annotations in the curated TAIR community database [46], the popular *ab initio* pipeline *MAKER* [44] predicted as few as 11% of the annotated *Arabidopsis* orphan genes, depending on the RNA-Seq evidence supplied [41].

Enhancing *ab initio* pipelines by other sequence-based information (e.g., motif/domain information, cellular location predictions, predicted isoelectric point (pI), genomics context) can improve gene predictions [47–49]. However, because it is not a given that newly evolved genes have canonical features, direct alignment of transcriptomic and/or proteomic data to the genome is critical for annotating orphan genes, as well as non-coding transcripts (lncRNAs, etc.) [3, 5, 7, 10, 32, 41, 48, 50].

Here, we reuse and re-mine aggregated RNA-Seq data to discover new potential gene candidates. The study comprehensively evaluates transcription and ribosomal binding of all open reading frames (ORFs) in the yeast genome over a wide variety of conditions, in the context of annotated genes. The research extends the results of previous studies, in that it globally represents ORFs in the *S. cerevisiae* genome across thousands of samples. Furthermore, we provide these data and extensive metadata via a biologist-friendly platform, MetaOmGraph (MOG [51], https://github.com/urmi-21/MetaOmGraph), which provides interactive, exploratory analysis [52] and visualization of expression levels, expression conditions, and co-expressed genes for the ORF-containing transcripts. This approach enables experimentalists to prioritize ORFs for functional characterization, and to logically define experimental parameters for these characterizations [51].

## Results

### Identifying potential cryptic orphan genes in *S. cerevisiae*

*S. cerevisiae* has the most extensively sequenced and annotated genome within the *Saccharomyces* genus, or perhaps across eukaryotes. However, despite the large body of research on *S. cerevisiae*, this genome expresses many transcripts not annotated as genes [3, 7, 9, 50, 53, 54], some of *de novo* origin [7, 26, 27, 39, 55], some supported with translational evidence [27, 50]. Our overall goal was to generate a comprehensive overview of expression of ORFs, and make this available in a format that can be readily explored. For this study, we classified all unannotated ORFs (>150 nt) and *Saccharomyces* genome database (SGD)-annotated genes in the *S. cerevisiae* genome according to phylostrata, transcription and translation evidence, and genomic context. We also included yeast ORFs < 150nt with transcription and/or translation evidence that had been characterized in two previous publications: smORFs [7] and txORFs [3]. Figure 1.B defines our terminology and lists the numbers of genes and ORFs in each category.

We inferred the oldest phylostratum (PS [56]) to which each *S. cerevisiae* protein (or candidate protein) could be traced, using the customizable *phylostratr* package [40] (Figure S1). Similarity to proteins of cellular organisms (i.e., proteins tracing back to prokaryotes) was designated as PS=1; no similarity to any protein outside of *S. cerevisiae* was designated as PS=15. (See supplementary file, *S.cerevisiae_RNA-seq_3457_27.mog* for full PS assignments by transcript). This analysis infers that fewer than 4% of SGD-annotated genes are orphans. In contrast, 54% of unannotated ORFs are orphans (“orphan-ORFs”), 40% are genus-specific (PS=10-14), and only 6% are more highly conserved (PS=1-9) (Figure 1.B).

In fungi, plants, and animals, the mean lengths of CDSs of annotated genes increase during evolution, with CDSs of orphan genes being the shortest [23, 27, 40, 57, 58] (Figure S2.A). The ORFs of yeast also follow a similar trend: average lengths of orphan-ORFs are shorter and average length of ORFs increases with increasing phylostrata (Figure S2). Consistent with the finding of Basile [59], the mean GC content for SGD-annotated orphan genes in *S. cerevisiae* is slightly lower (though not statistically significant) than that of more conserved genes. Like the SGD-annotated orphan genes, the Q3-transcribed orphan-ORFs (ORFs in top quartile of mean expression, see Figure 1.B) have a slightly lower mean GC content than ORFs of other phylostratum levels (Figure S2.B). Vakirlis [55] reported a higher mean GC content among those orphan genes that have a confirmed *de novo* origin.

### Transcriptional landscape of genes and ORFs

Expression of many annotated orphan genes is developmentally localized, up-regulated under environmental stress, or associated with species-specific traits [23, 60–63]. For example, more yeast orphans are ribosomally-bound under starvation conditions than control conditions [6, 7]. We anticipated that sparse-expression would be a characteristic of many of those orphan-ORFs that are actually orphan *genes* that have escaped annotation. To capture expression of these orphan-ORFs, we deemed it essential to use RNA-Seq samples comprising diverse developmental, genetic, and environmental conditions.

RNA-Seq samples drawn from a wide range of conditions have an added benefit. Because orphans have no homologs in other species, and no recognizable functional domains, these characteristics cannot be used to provide a clue as to function [23], rendering functional inference a particular challenge. The assumption that genes with similar patterns of expression are likely to encode proteins involved in a common process provides a powerful approach to infer experimentally-testable functions for genes of unknown function. Therefore, using datasets incorporating the diverse conditions in which orphans-ORFs or orphan genes might be expressed is key to functional inference and to determine the conditions that induce their expression.

To gather RNA-Seq data from diverse conditions, we collected raw sequence reads and metadata of 3,457 RNA-Seq samples from 177 studies in The National Center for Biotechnology Information-Sequence Read Archive (NCBI-SRA). (See *S.cerevisiae_RNA-seq_3457_27.mog* for metadata and counts). The experimental variables across these samples include a wide variety of mutants, chemical treatments, stresses, nutrition deprivations, and growth stages. We quantified the expression of all 29,354 ORFs and 6,692 SGD-annotated genes of *S. cerevisiae* across the 3,457 RNA-Seq samples.

Figure 2 shows a heatmap for expression of SGD-annotated genes, smORFs (sequences encoding small orphan proteins with ribosomal evidence of translation [7]), and transcribed orphan-ORFs (> 150 nt) across the 3,457 RNA-Seq samples. (See Figure S3 for expression plot of all genes and ORFs). The mean expression across all samples for SGD-annotated genes is 38 cpm, whereas the mean expression for the Q3-transcribed ORFs is 18 cpm (Table S1). Many SGD-annotated genes are expressed in most of the samples. In contrast, as we anticipated based on the erratic pattern of expression of annotated orphan genes, most of the orphan-ORFs show very low expression in most RNA-Seq samples, but accumulate highly in a few samples. This sporadic expression contributes significantly to the observed lower mean expression of the orphans. It also demonstrates how many transcribed sequences might be missed if smaller, less diverse datasets are analysed.

**Figure 2.**
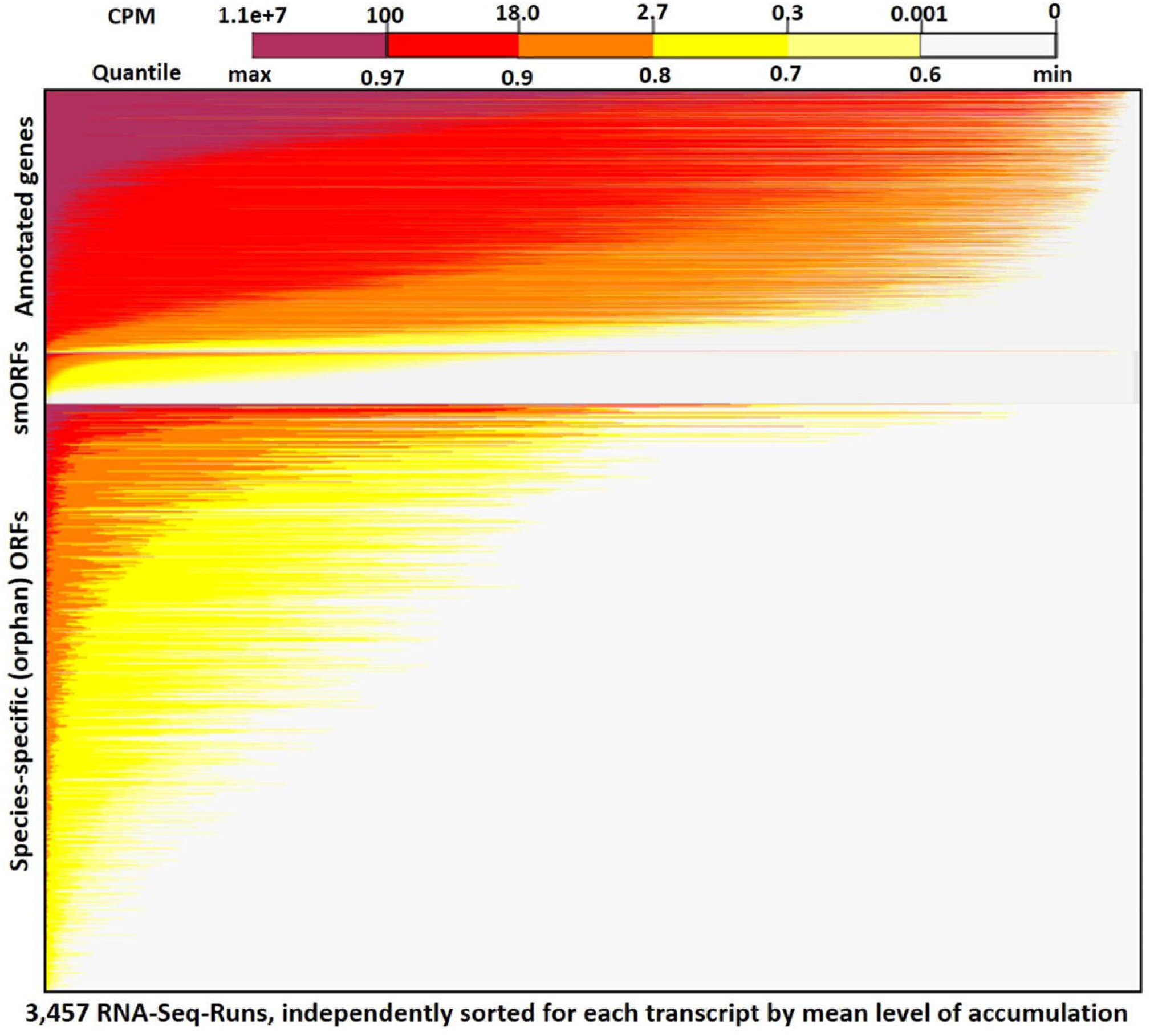
RNA-Seq expression heatmap across 3,457 samples for orphan-ORFs and SGD-annotated genes. Top panel, SGD-annotated genes (6,692); middle panel, smORFs (Carvones et al., 2012) (1,139); bottom panel, orphan-ORFs (15,805). (See Figure S3 for all transcript classes). Each row represents a transcript. Within a panel, each transcript is ordered by its mean cpm. Within each row, the 3,457 samples are sorted independently by highest expression of the transcript. The restricted conditions of expression of many orphan-ORFs is visually apparent.

Ninety-nine percent of the 3,457 RNA-Seq samples have transcription evidence for at least one of the Q3-transcribed ORFs (Figure 3). Some samples are particularly rich in Q3-transcribed ORFs. For example, 50 samples have transcription evidence for >1,200 of the Q3-transcribed ORFs; 47 of these samples are from wild type strains, many grown under conditions of nutritional or chemical stress.

**Figure 3.**
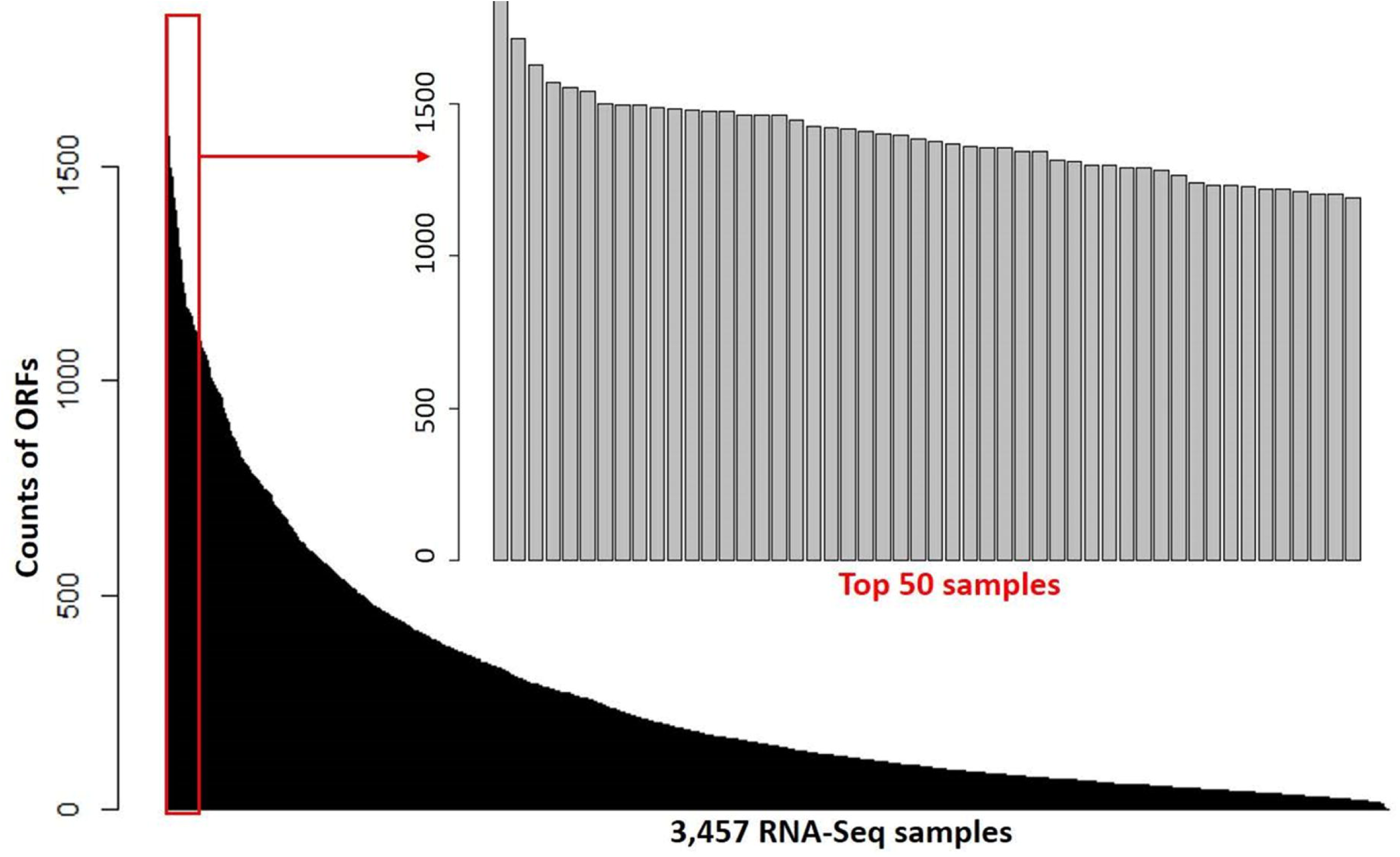
Counts of highly-transcribed (Q3) ORFs in each RNA-Seq sample. The black bars show distribution of the counts of the 8193 Q3-transcribed ORFs. X-axis, 3,457 RNA-Seq samples, sorted by counts. The grey bar inset details the 50 RNA-Seq samples with the largest number of Q3-transcribed ORFs; each of these samples contains over 1200 Q3-transcribed ORFs.

The conserved SGD-annotated genes have higher mean expression than either the orphan SGD-annotated genes, the Q3-transcribed orphan-ORFs, or the Q3-transcribed conserved-ORFs (Kolmogorov-Smirnov Test, p-values < 0.001; Figure 4). However, over 600 orphan-ORFs have a higher mean expression than 10% of conserved SGD-annotated genes, 289 orphan-ORFs have a mean expression higher than 25% of the conserved SGD-annotated genes, and 36 orphan-ORFs have a mean expression higher than 90% of conserved SGD-annotated genes (Figure 4 and Table S1. A).

**Figure 4.**
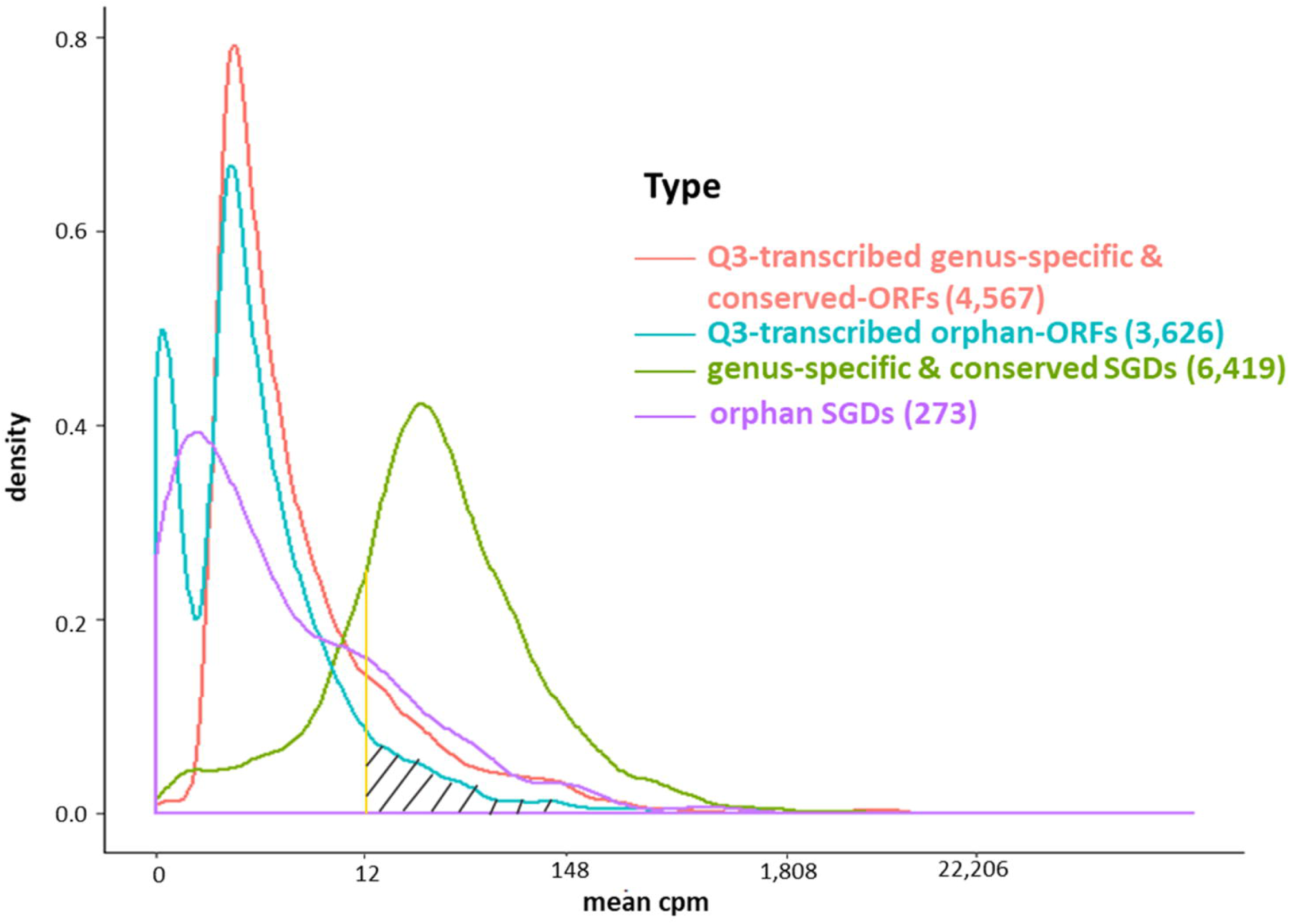
Density plot of mean expression level of transcripts across 3,457 samples for SGD-annotated genes and Q3-transcribed ORFs. X-axis, *edgeR*-normalized mean expression of genes and ORFs. Y-axis, number of transcripts. The area under the curve of the density function represents the probability of a range of mean cpm. The bimodal curve of all orphan-ORFs is attributable to the low mean expression of the smORFs (see Figure S4, S5). About half of the Q3-transcribed orphan-ORFs have higher mean expression than orphan SGD-annotated genes. Over 600 orphan-ORFs have a higher mean expression than 10% of conserved SGD-annotated genes; 289 orphan-ORFs (gray hatched area) have a higher mean expression than 25% of conserved SGD-annotated genes; and, 36 orphan-ORFs have a mean expression higher than 90% of conserved SGD-annotated genes (See also Table S2)

### Genomic context of the ORFs

We surveyed the genomic location of each ORF relative to the nearest SGD-annotated gene (Figure S6). A recent study using on two experimental conditions [50] reported that a high proportion of expressed but unannotated transcripts in yeast overlap known CDSs but are transcribed from the opposite strand. Consistent with [50], 30% of the transcribed ORFs overlap CDSs and are transcribed from the opposite strand (reverse orientation) in our study. Furthermore, regardless of orientation (same *versus* reverse), ORFs that overlap an annotated CDS have a median of mean expression *5-fold higher* than ORFs located within or outside a CDS (Wilcoxon rank-sum test, p-value < 0.001, Figure S6). We have no explanation for this phenomenon. The orientation in which overlapping ORFs are transcribed relative to the associated CDSs does not significantly alter the mean expression level of the ORFs (Figure S6).

Of the 289 orphan-ORFs with the highest transcription (Figure 4), 49% overlap an annotated CDS, rather than being within an annotated CDS or outside an annotated CDS (Figure S7). This is significantly higher (Fisher’s exact test, p-value< 0.001) than the proportion among all ORFs (15%) and all orphan-ORFs (14%) that overlap CDS (Figures S6, S7). Most of the 289 orphan-ORFs that overlap a CDS have a reverse orientation (convergent or divergent) relative to the SGD-annotated gene they overlap (Figure S7).

Many RNAs in fungi and humans that have been annotated as “lncRNAs” are associated with ribosomes, and/or have proteomics evidence, indicating some of them may function as protein-coding genes [2, 6, 11, 32, 64]. To examine translation evidence in our study, we globally evaluated translation evidence, mapping raw reads from 302 ribosomal profiling RNA-Seq (Ribo-Seq) samples in SRA to the unannotated ORFs and SGD-annotated genes of *S. cerevisiae.* (See supplementary file *Ribo-Seq_counts.csv* and *Ribo-Seq_metadata.xlsx* for raw counts and metadata). About 61% of Q3-transcribed conserved-ORFs, 40% of genus-specific-ORFs, and 51% of orphan-ORFs have translational evidence among these Ribo-Seq samples (Figure 1.B). This compares to 97% of the conserved SGD-annotated genes, 45% of genus-specific SGD-annotated genes, and 38% of orphan SGD-annotated genes. The mean Ribo-Seq raw counts were significantly different (t-test p-value < 0.001) among classes of transcripts, depending on whether they were orphan, genus-specific, or conserved (Figure 5.A). The mean Ribo-Seq raw counts for the low-transcribed ORFs are significantly lower than for the Q3-transcribed ORFs, and the mean Ribo-Seq raw counts for the ORFs with no transcription evidence are 0 or near 0 (Figure S8).

**Figure 5.**
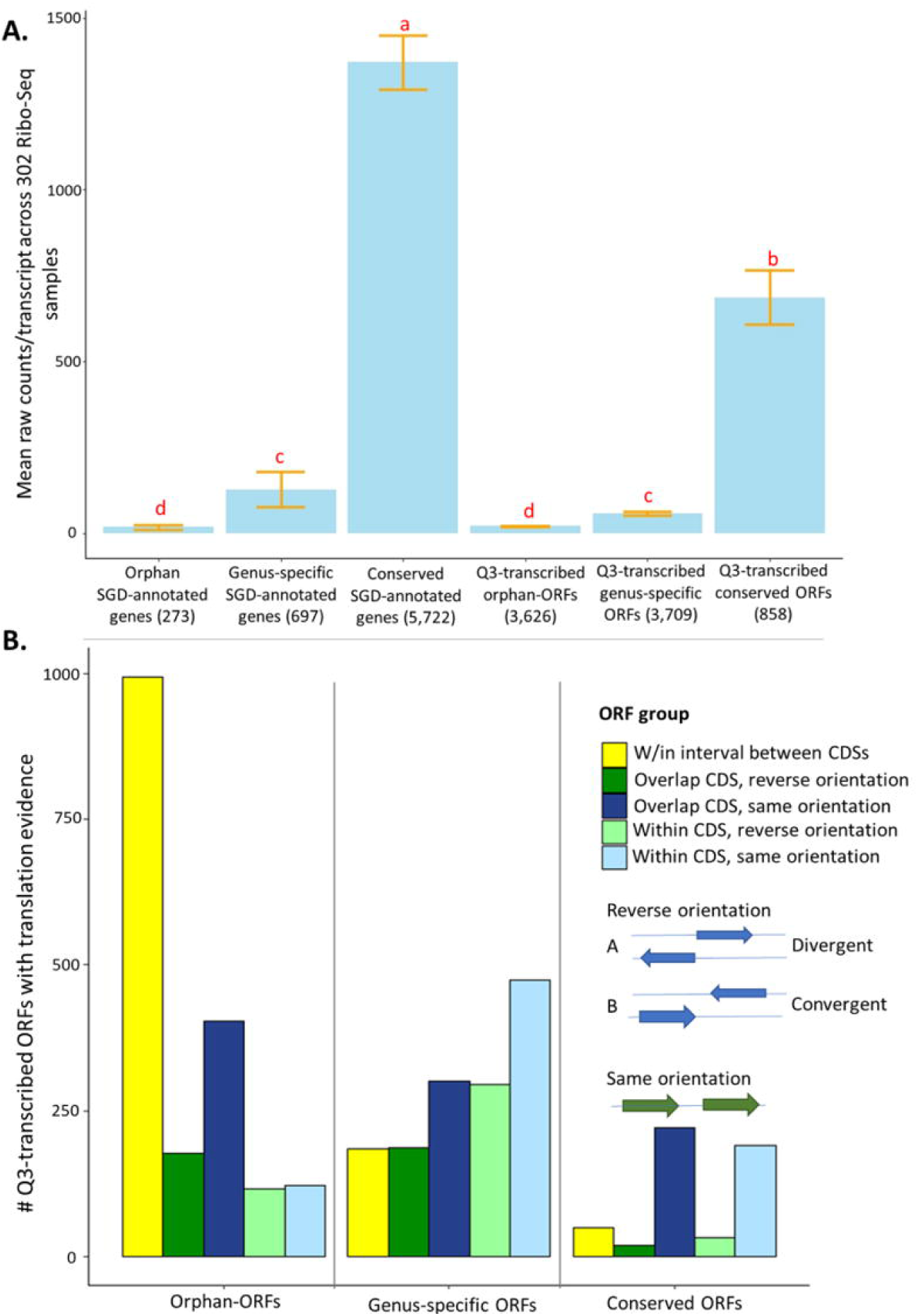
Mean expression and numbers of genes and ORFs with translational evidence, partitioned by phylostrata and genomic context. Ribo-Seq data were analysed for genes and ORFs across 302 samples using *ribotricer* [15]. **A.** Mean raw Ribo-Seq counts/transcript for all genes and ORFs. X-axis, genes and ORFs as classified by phylostrata. Y-axis, mean raw counts. The letters above each bar indicate significance in each group according to a t-test (p-value cutoff is 0.01). Similar to mean RNA-Seq counts, the conserved genes and conserved-ORFs have more total mean Ribo-Seq counts. **B.** The 3,857 Q3-transcribed ORFs that had Ribo-Seq translation evidence were divided into groups according to their relationship to annotated CDS (see also Figure S6), and the numbers of genes and ORFs with translational evidence was determined. The gene/ORF with mean counts across 302 Ribo-Seq samples higher than 0.3 was consider to have translation evidence. X-axis, groups of genes and ORFs, classified by phylostrata. Y-axis, number of ORFs in each group. The proportions of ORFs are significantly different among three groups according to a chi-square test (p-value<0.001). Over half the orphan-ORFs with translation evidence are located between CDSs.

The proportions of Q3-transcribed ORFs with translation evidence located within, overlapping, or between annotated CDSs are significantly different among orphan-ORFs, genus-specific-ORFs, and conserved-ORFs (Chi-square test, p-value < 0.001) (Figure 5.B). Notably, 54% of Q3-transcribed orphan-ORFs with translation evidence are located in the intervals between annotated CDSs, compared to only 12% of the genus-specific ORFs and 9% of the conserved ORFs (Figure 5.B).

Since yeast was the first model eukaryotic genome [65], and has been reannotated over time, it would be expected that most conserved genes are already annotated. However, some genus-specific-genes might have been missed because homology is a major criterion used for genome annotation. Orphan genes, which have no homologs in other species, sparser expression, and likely fewer canonical features [41], are yet less likely to have been annotated. In total, 1,477 Q3-transcribed genus-specific-ORFs and 1,850 Q3-transcribed orphan-ORFs have ribosomal binding evidence. These transcribed, translated ORFs are candidates as protein-coding genes.

Five hundred and thirty of the 858 Q3-transcribed conserved-ORFs also have translation evidence. There are several possible explanations for why a transcript with homologs in other species are not annotated as genes. Some of these conserved-ORFs may be pseudogenes that retain some homology and expression, but have lost functional capacity. Other conserved-ORFs might encode active proteins, by because they are expressed only under limited conditions they might not have been sampled when SGD annotations were made. Other conserved-ORFs may have been ignored because their ORF codes for a shorter protein than the canonical gene family member. (On average, a Q3-transcribed conserved-ORF is significantly shorter than the homologous SGD-annotated gene (t-test, p-value < 0.001)). However, it not a given that because an ORF encodes a shorter protein it is non-functional. Shorter homologs of proteins with known function may play a biological role in regulating signal transduction, modulating enzyme activity, and/or affecting protein complexes, potentially competing with their “full-length” homolog [66, 67]. Translation of a short conserved-ORF also might be regulatory, in that it limits translation of a nearby active protein [68].

### Network inference and co-expression analysis

To analyse the expression patterns of the ORFs in the context of annotated genes, we optimized correlation and network parameters for the RNA-Seq expression data (see Methods, Figures S10, S11, and Table S3), and focused our subsequent interactive co-expression analysis and visualization on a dataset (“SGD+ORF” dataset) composed of 14,885 transcripts (all SGD-annotated genes; the 7,054 Q3-transcribed ORFs; and all 1139 smORFs) across 3,457 RNA-Seq samples.

We then computed the Pearson pairwise correlation (PCC) matrix for the SGD+ORF dataset, and partitioned the resultant PCC matrix by Markov chain graph clustering (MCL) [69] into 544 clusters (Table S4 for overview; genes and ORFs with cluster designations at supplementary file *S.cerevisiae_RNA-seq.mog*). Forty-six percent of the 273 SGD-annotated orphan genes and 59% of the 3,899 Q3-transcribed orphan-ORFs are members of clusters containing more than five genes and include genes of known function, thus providing potential for functional inference.

It was possible that ORF expression might be correlated with that of adjacent or overlapping SGD-annotated genes, i.e., that ORFs are expressed due to a physical proximity to transcribed SGD-annotated genes. We used two approaches to evaluate the extent to which such “piggybacking” might occur. In the first approach, we focused on the 390 ORFs that are located completely within UTRs of SGD-annotated genes (88% are orphan-ORFs). About 80% of these ORFs have a PCC less than 0.6 (0.6 is the correlation cut-off we used for MCL) with the encompassing SGD-annotated genes, however, about 2% (eight) ORFs have a correlation higher than 0.9. In the second approach, we calculated how many ORFs are in the same cluster as nearby annotated genes. To do this, we randomly selected 366 ORFs that were members of clusters, and made test clusters of the same sizes, each cluster containing randomly-selected SGD-annotated genes and the identical ORFs as in the experimental data. Then, we calculated the distance of each ORF to each SGD-annotated gene in the randomly-created and the experimental clusters. The distances were not statistically different in the experimental versus the random clusters (p-value=0.16 in a t-test for difference). These tests indicate that the expression of ORFs is not generally associated with the ORFs being within or near to an SGD-annotated gene, and co-expressed with it. However, there is strong support for such a relationship in specific cases (e.g., Figure 9, and as reported in [55]).

About 65% of the Q3-transcribed ORFs are assigned to clusters in the co-expression matrix. Regardless of whether they are protein-coding, they could play a biological role. The highly transcribed ORFs with translational activity provide an evidence-based cadre of *candidate* protein-coding genes that could be experimentally tested.

### GO enrichment analysis for co-expressed clusters

In order to evaluate the significance of the clustering results, we compared the extent of enrichment of Gene Ontology (GO) terms in the set of clusters obtained from MCL-partitioning experimental data to that of 100 randomly-generated sets of clusters. For each randomly-generated set, the number of clusters and the number of genes per cluster were held the same as the set of clusters from the experimental data; however, the genes assigned to each cluster were changed by random permutation. The best adjusted p-value for enriched GO terms was recorded for each cluster and averaged across all clusters to obtain a mean best p-value [70] (Figure 6). Distribution of the p-values for GO terms in the 100 sets of randomized clusters was compared to that of the experimental data (red arrows in Figure 6). For each GO ontology category (Biological Process (BP), Cellular Component (CC), and Molecular Function (MF)), the best mean p-values for the experimental data are 0.019, 0.023, and 0.027, respectively. These values are significantly better than those of any of the randomly-obtained cluster sets, indicating that the MCL gene clusters derived from the experimental data is not random. Co-expressed genes are implicated as being involved in a similar process [71, 72]. That this study is based on over 3,000 biological conditions further strengthens the likelihood that genes in each cluster might share a related biological process.

**Figure 6.**
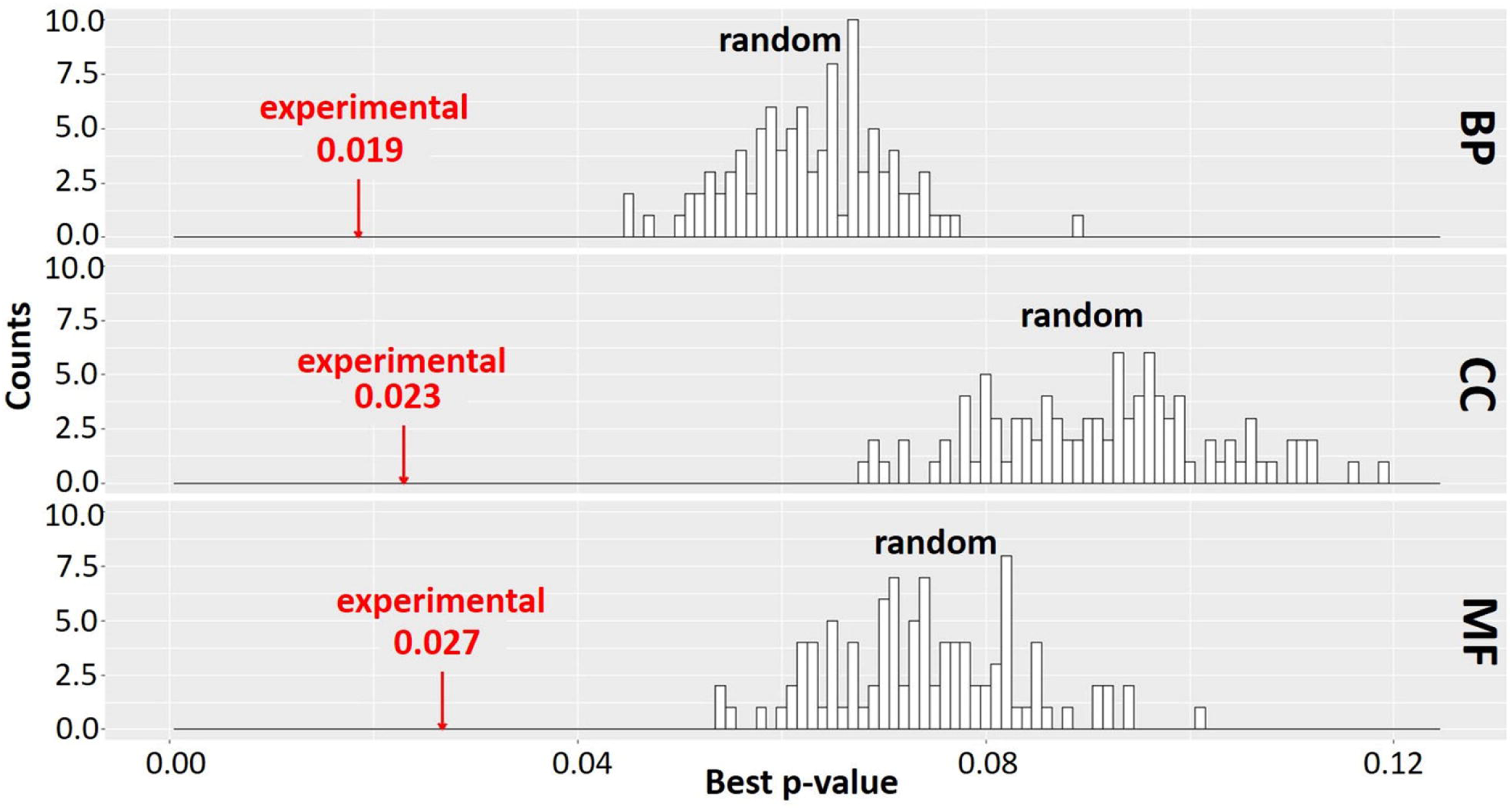
GO enrichment analysis of experimental data and random test distribution. A Pearson correlation matrix of the SGD+ORF dataset was partitioned into clusters by MCL. Best p-values (mean of the lowest adjusted p-values for GO terms) were determined across all clusters of the experimental data and all clusters of random permutations, similar to [70] (Section 7 in *Supplementary_Material.pdf*). Red arrow, experimental data. Black bars, best p-value of 100 randomly-obtained permutations with size and number of clusters identical to experimental data. BP, biological process; CC, cellular component; MF, molecular function. The clustering result is significantly better for experimental data than any random permutation.

### Exploring Gene Function: Case study, Cluster 112

The co-expression clusters are often composed of genes and ORFs distributed across spatially diverse regions of the genome (For a list of all genes and ORFs as partitioned into clusters by MCL, see supplementary file *S.cerevisiae_RNA-seq_3457_27.mog)*. For example, MCL Cluster 112 (Figure 7) contains 20 SGD-annotated genes and 21 unannotated ORFs dispersed on 14 chromosomes. Twelve of the genes are in the seripauperin (*PAU*) family. The molecular function of the *PAU* genes is not known. However, *PAU*-rich co-expressed gene clusters have been identified in independent microarray studies [73, 74]. Many *PAU*s are induced by low temperature and anaerobic conditions, and repressed by heme (Rachidi, Martinez, Barre & Blondin, 2000) and individual *PAU* proteins confer resistances to biotic and abiotic stresses [76]. *YER011W* and *YJR150C*, also in Cluster 112, are localized to the same cellular compartments as *PAU*s and are also induced under anaerobic conditions [77–80]. The other SGD-annotated genes in this cluster have no functional description. GO enrichment analysis identified eight GO terms as significantly-over-represented in Cluster 112 (Table 1). Figure 8 represents a case study of an approach to develop a meaningful hypothesis. The example shows the expression of the genes and ORFs in Cluster 112 across all 3545 samples of the RNA-Seq SGD+ORF dataset, and highlights the two studies that evaluate oxygen content as an experimental variable. Study SRP067275 compares four growth stages of the stress-tolerant yeast strain GLBRCY22-3 grown in YPDX and ACSH media, with and without oxygen [81] (Figure 8, top left); the expression of the genes and ORFs in Cluster 112 is higher under anaerobic conditions, irrespective of media or growth stage. Study SRP098655 compares *OLE1*-repressible strains growing under anaerobic and aerobic conditions [82] (Figure 8, top right); expression of genes and ORFs in Cluster 112 is induced in cells grown under anaerobic conditions. These expression patterns indicate the genes and the ORFs in this cluster might be sensitive to anoxia, or might play a role in cellular response to this stress.

**Table 1.** Significantly-enriched GO terms in Cluster 112. Based on the GO terms assigned to the gene members of known function, Cluster 112 is enriched in the GO terms shown in Table 1. The results indicate a possible role in stress response related to the cell wall for the ORF members of Cluster 112. (See *S. cerevisiae_RNA-seq_3457_27.mog* for complete clustering and ontology results).

| Ontology | GO name | Adjust p-value |
| --- | --- | --- |
| MF | structural constituent of cell wall | 1.90E-27 |
| BP | response to stress | 3.51E-27 |
| CC | fungus-type cell wall | 1.62E-20 |
| BP | fungus-type cell wall organization | 1.34E-19 |
| CC | fungus-type vacuole | 3.60E-05 |
| CC | cell wall | 1.99E-03 |
| CC | extracellular region | 6.18E-03 |
| CC | anchored component of membrane | 1.87E-02 |

**Figure 7.**
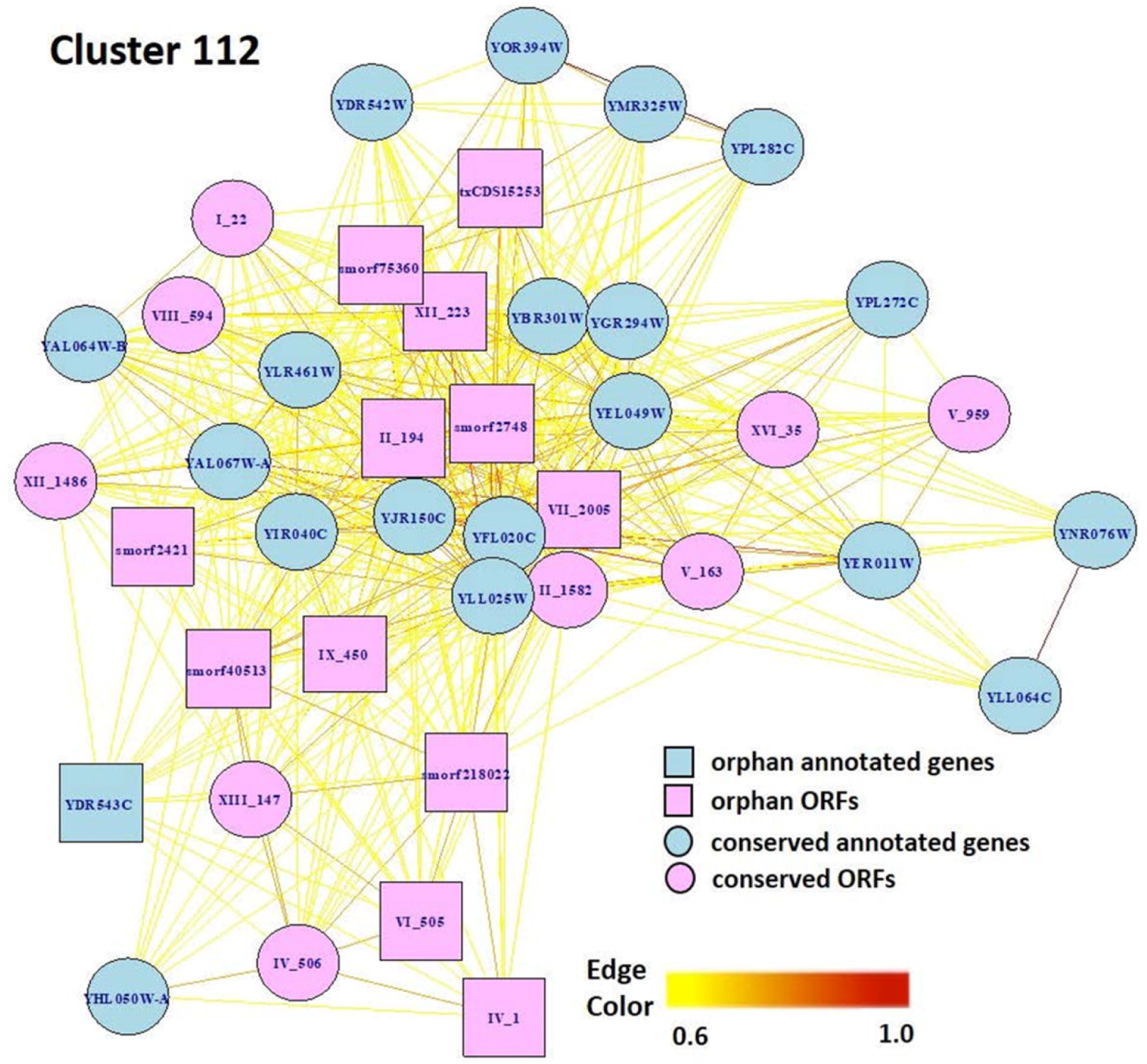
Network view of genes and ORFs in Cluster 112. A Pearson correlation matrix of the SGD+ORF dataset was partitioned into clusters by MCL. Cluster 112 is an example of a cluster containing SGD-annotated genes and ORFs, including orphans. Edge colors, Pearson correlations of 0.6 to 1.0. Visualization by *igraph* in R [99].

**Figure 8.**
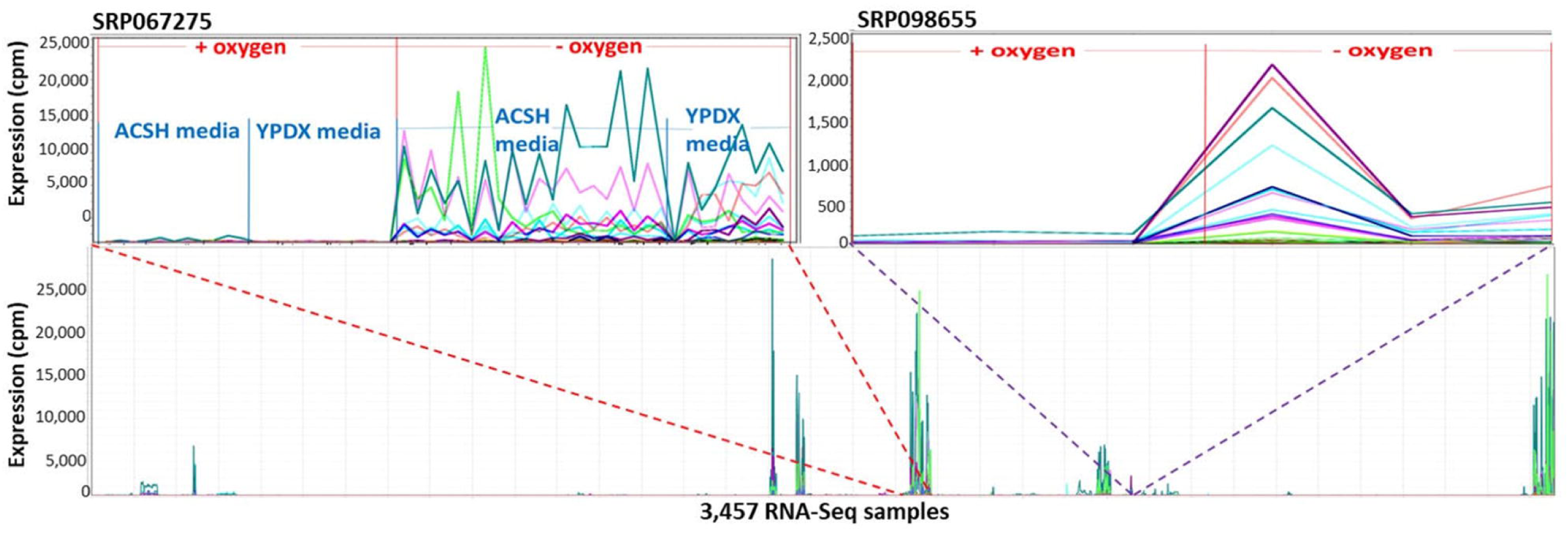
The 41 genes and ORFs in Cluster 112 respond to anoxia. A Pearson correlation matrix of the SGD+ORF dataset was partitioned into clusters by MCL. The 41 genes and ORFs in Cluster 112 are co-expressed across multiple conditions. X-axis, 3,457-samples, sorted by study. Y-axis, expression values. Each line represents the expression pattern of a single gene or ORF. Top left inset, zoom-in to visualize Study SRP067275. RNA-Seq samples sorted by: aerobic or anaerobic condition, ACSH or YPDX media, and growth phase. ACSH, Ammonia Fiber Expansion-(AFEX-) pretreated corn stover hydrolysate. YPDX, YP media containing 60 g/L and 30 g/L xylose. Top right inset, zoom-in to visualize Study SRP098655. The genes and ORFs are up-regulated in response to anoxia, regardless of changes in growth media. No ORF in Cluster 112 is located near an SGD-annotated gene in Cluster 112. Visualizations and co-expression calculations by MOG (Singh et al., 2020).

### Exploring Gene Function: Case study, *smORF247301*

Though rare, some transcribed ORFs that are located near or in an existing gene share a similar transcription pattern. An example is *smORF247301*, one of the most highly expressed smORFs, which is 77 nt upstream of *YPL223C* (Figure 9). MOG analysis indicates *smORF247301* and the SGD-annotated gene *YPL223C* have a PCC of 0.95 across the 3,457 RNA-Seq samples. *smORF247301* is located on the “+” strand of chromosome 16, while *YPL223C* is on the “-” strand of the same chromosome. The CDS of *YPL223C* is 507 nt, while *smORF247301* is 33 nt. *YPL223C* is more highly expressed than *smORF247301. YPL223C*, a hydrophilin gene that is essential in surviving desiccation-rehydration, is regulated by the high-osmolarity glycerol (HOG) pathway [83], and induced by osmotic, ionic, oxidative, heat shock and heavy metals stresses. Analysis using MOG shows *smORF247301* and *YPL223C* have increased expression in response to osmotic, heat, and desiccation stresses in three independent studies (Figure 9 B-D). *smORF247301* has translation evidence ([7] and this study).

**Figure 9.**
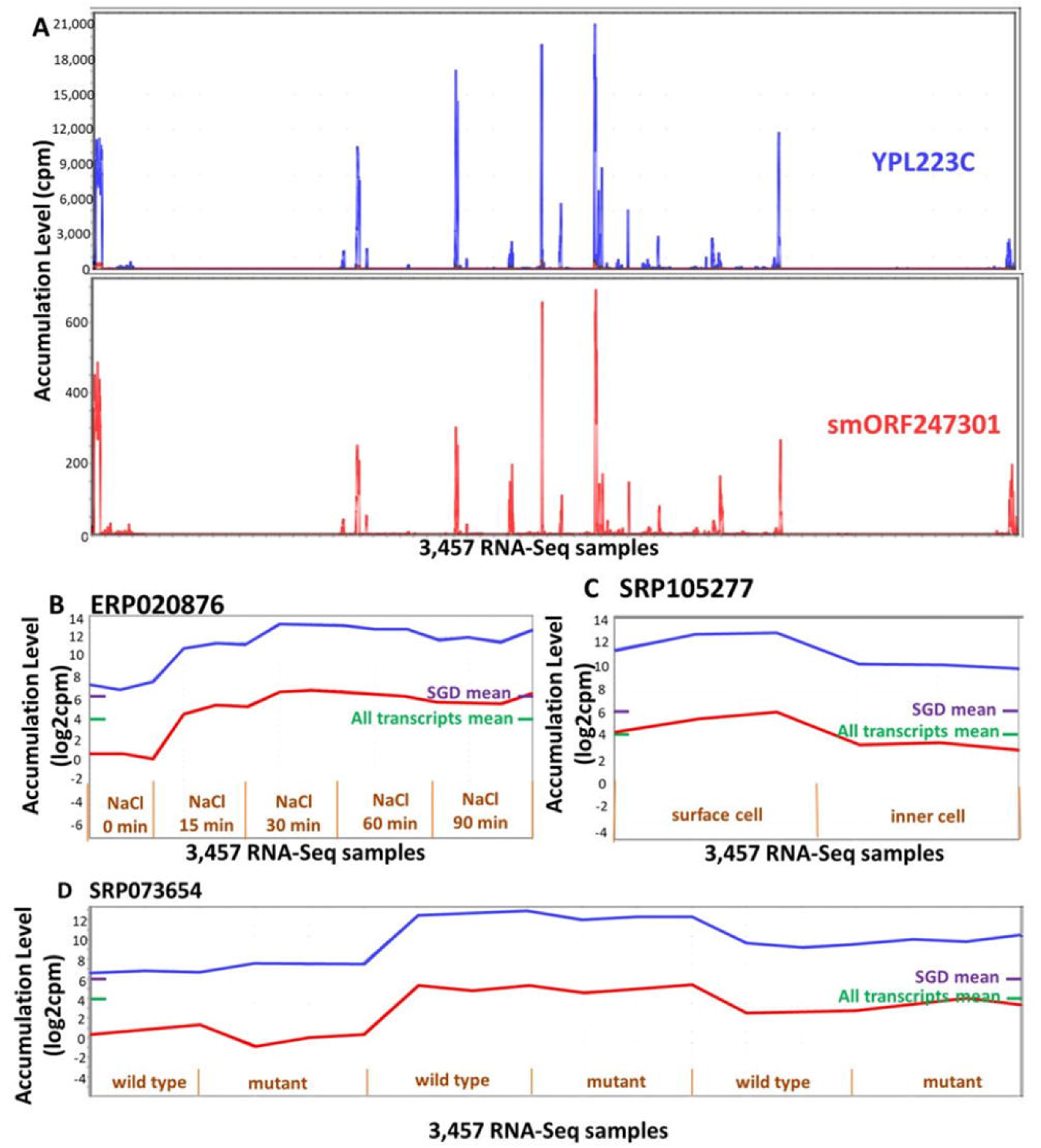
Expression patterns of smORF247301 and YPL223C. *smORF247301* and *YPL223C* are located on adjacent regions of chromosome 16 and are transcribed in convergent orientation. **A.** Expression patterns are similar (Pearson correlation, 0.95) across 3,457 samples. **B-D.** Expression patterns for smORF247301, YPL223C in three studies. X-axis, 3 samples per treatment. Purple bar on right side of panels, mean expression level of all SGD-annotated genes; Green bar on right side of panels, mean expression level of all SGD-annotated genes plus ORFs. Visualizations and co-expression calculations produced by MOG (Singh et al., 2020).

It is possible that the transcription and translation of *smORF247301* is “noise” (Eling, Morgan & Marioni, 2019) associated with the expression of the nearby *YPL223C*. A second possibility is that *smORF247301* is a young, not-yet-annotated gene. It might be “piggybacking” on the expression apparatus of *YPL223C*. However, *smORF247301* and *YPL223C* are transcribed in a *convergent* orientation (-> <-, Figure 5B); thus, the process, described by [55], whereby two transcripts in divergent orientation (<- ->, Figure 5B) are co-expressed via a common bidirectional promoter would not apply in the case of *smORF247301* and *YPL223C*. A different “piggybacking” mechanism might apply: perhaps, due to its location in open chromatin, *smORF247301* is provided with a ready-made exposure to transcription factors when gene *YPL223C* is transcribed. If a transcript (e.g., *smORF247301*) conferred a survival advantage under the same conditions as did its established neighbouring gene (e.g., *YPL223C*), it could emerge as a new, co-expressed, gene by this mechanism.

Five hundred and thirty-seven orphan-ORFs with transcription and translation evidence are in physical proximity to an SGD-annotated gene and are transcribed in a *divergent* orientation (see supplementary file, *divergent_pairs.csv*). Of these pairs, 12 are co-expressed (PCC > 0.6); these 12 ORFs are potentially co-expressed by a bidirectional promoter (e.g., as described by [55]) The 525 orphan-ORFs that are not co-expressed, might still be controlled by a bidirectional promoter, because yeast ORFs can be transcribed by a bidirectional promoter, but not be correlated in expression because they are influenced by different transcription factors [85].

### Future studies

The SGD+ORF dataset we provide can be reanalysed by different approaches. Each combination of network inference and partitioning approaches can supply complementary information. For example, networks can be inferred by correlation, mutual information [86], or relatedness approaches [87]. Pearson correlation, used here, is highly sensitive at extracting genes whose expression is linearly correlated across multiple conditions, but misses non-linear co-expression. Likewise, networks can be partitioned by several methods, e.g., MCL (as in this study), Modularity [88], or a promising new approach, Reduced Network Extreme Ensemble Learning (RenEEL) [89]. There has been little investigation into the strengths and weaknesses of the various inference and partitioning methods for extracting different types of biological information.

Moreover, we focus here on protein-coding transcripts; similar investigations using diverse RNA-Seq data could center on non-coding RNAs or transcript-encoding very small proteins.

The information resulting from such studies can easily be incorporated into a new MOG project to enable interactive analysis and visualization.

## Conclusion

In this study we have globally assessed the accumulation of transcripts representing 36,046 annotated genes and unannotated ORFs of *S. cerevisiae* across 3,457 public RNA-Seq samples derived from diverse biological conditions. Ninety-five per cent of the transcribed ORFs are orphans or genus-specific. Despite a strong tendency to be transcribed only under restricted conditions, 269 orphan-ORFs had mean levels of transcription greater than 25% of SGD-annotated genes. Over 2,000 transcribed ORFs with translation evidence are members of co-expression clusters, providing additional clues as to a potential function.

The proportion of transcribed and translated ORFs that are functional is completely unknown. The SGD+ORF dataset assembled herein represents expression of SGD-annotated genes and unannotated ORFs under multiple conditions; it is delivered in a readily explorable, user-friendly format via the MOG platform. Combining this network-informed view of aggregate RNA-Seq data with text-mining of sample and gene metadata creates a powerful approach to develop novel, experimentally-testable hypotheses on the potential functions of as-yet-unannotated transcripts.

## Materials and methods

### Extracting ORFs and delineating orphan-ORFs in *S. cerevisiae*

ORFs (>150 nt) that were not annotated in SGD as CDS, were extracted from the yeast genome (version: R64-1-1) by bedtools2 [90], and translated by emboss [91], yielding 24,912 ORFs. To these ORFs we added two sets of ORFs <150 nt identified in other studies: the 1,139 small translated sequences (smORFs) identified by ribosome profiling [7] and the 3,303 of ORFs identified by TIF-Seq (txCDS) [3] that were less than 150 nt (thus, not included in the bedtools2 extraction). These 29,354 ORFs, together with the 6,692 protein-coding genes annotated in SGD, were subjected to phylostratigraphic analysis.

We inferred the phylostratum for 29,354 ORFs and 6,692 SGD-annotated protein-coding genes via the R package, *phylostratr* [40]. The analysis compared the proteins predicted from the *S. cerevisiae* ORFs to proteins of 123 target species distributed across phylostrata: 117 species identified by the *phylostratr* algorithm, supplemented with six manually-selected species in the Saccharomyces genus (*S. paradoxus, S. mikatae, S. kudriavzevii, S. arboricola, S. eubayanus*, and *S. uvarum*). To minimize false positives when identifying orphan ORFs and CDS from *S. cerevisiae*, we took advantage of the customization capabilities of *phylostratr* and included the predicted translation products from all ORFs (>150 nt) from each of the six *Saccharomyces* genomes, in addition to all SGD-annotated proteins of these species. (See *Supplementary_Material.pdf*, Figure S1 for workflow, Section 13 for full species list, and *phylostratr_heatmap.pdf* for gene by gene (and ORF by ORF) heatmap). Each gene was assigned to the most evolutionarily-distant phylostratum that contains an inferred homolog. A gene or ORF is inferred to be an orphan if its encoded protein is assigned the phylostratum level *S. cerevisiae*. A BLASTP for each ORF and CDS in *S. cerevisiae* against *Saccharomyces* spp ORFs and CDS gave identical results to those of *phylostratr* in identifying the orphan genes and ORFs.

### Raw read processing and network optimization

Our RNA-Seq data analysis pipeline is shown in Figure S9. We selected all samples with *S. cerevisiae* taxon ID 4932, Illumina platform, and paired layout from NCBI-SRA and then filtered out samples with miRNA-Seq, ncRNA-Seq, or RIP-Seq library strategies. In total, we collected raw reads data (FASTQ format) and metadata from 3,457 RNA-Seq samples (177 studies). A transcriptome was created from SGD-annotated cDNA and unannotated ORFs, and then expression levels of annotated genes and ORFs over the 3,457 RNA-Seq samples were quantified by *kallisto* [92] (See supplementary file *S.cerevisiae_RNA-seq.mog* for RNA-Seq metadata and normalized cpm data; all data including raw counts is accessible at DataHub (https://datahub.io/lijing28101/yeast_supplementary)).

We evaluated the performance of two diverse normalization methods for the raw count data (Section 8 in *Supplementary_Material.pdf*). We normalized raw counts by *edgeR* [93] based on the evaluation of [94]. We also normalized the same data by a single cell RNA-Seq normalization approach *SCnorm* [95]. This method examines sequence information from individual cells with the aim to provide a higher resolution of cellular differences. We tested this method because two features of single cell RNA-Seq data are similar to the orphan-ORF-focused multi-study data assembled for our study: 1) raw counts contain an abundance of zero-expression values, 2) technical variability among samples is high. After normalization, only ORFs with mean expression values in the upper quantile of mean expression (Q3-transcribed) were retained. We generated two datasets: 1) all SGD-annotated genes (SGD dataset); and 2) all Q3-transcribed ORFs, smORFs, and SGD-annotated genes (SGD+ORF dataset). For each normalization approach and dataset, we calculated pairwise Pearson correlation matrices among all 3,457 RNA-Seq samples.

Three PCC cutoffs (0.6, 0.7 and 0.8) were used to create networks of different densities from the matrices (Section 7 in *Supplementary_Material.pdf*). We then applied MCL to partition each network using our in-house Java Spark implementation (GitHub: https://github.com/lijing28101/SPARK_MCL) designed to optimize efficiency. All data analysis in this work, except for MCL clustering and RNA-Seq expression visualization, were performed in R software.

### Cluster evaluation by GO term enrichment analysis

Clusters resulting from each of the eight MCL analyses obtained from the different normalization methods and PCCs were evaluated by GO enrichment analysis using *clusterProfiler* [96]; in this evaluation, only clusters with over five genes were considered (Section 7 in *Supplementary_Material.pdf*). The GO term enrichment of each experimental result was compared to that of 100 random sets of clusters, which were obtained by permuting gene IDs. For these permutations, the same number of clusters of the same size as those from the experimental result were assigned to each random set using the method of [70]. The best adjusted p-value (pmin, smallest adjusted p-value) was recorded for the enriched GO terms in each cluster. Each random cluster set was assigned a score Si, which is the average pmin across all clusters in the set.

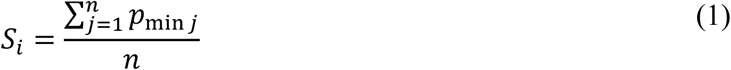

where n indicates the number of clusters. The distribution of S values for GO classes, Biological Process, Cellular Component, and Molecular Function, for random sets were compared to the respective values for the real experimental data. In each ontology, the experimental score was less than any of the random scores, indicating that experimental data have biological significance (permutation test, p-value=0). Based on the GO enrichment results we chose *edgeR* normalization (Section 8 in *Supplementary_Material.pdf*) and a PCC of 0.6 (Section 7 in *Supplementary_Material.pdf*) for future analyses.

### Ribo-Seq analysis

To investigate the translational activity of unannotated ORFs, we analysed 302 samples (23 studies) of yeast Ribo-Seq data; this represented about half of the available Ribo-Seq in the SRA database. Raw reads (SRA-formatted) were downloaded, and the SRA toolkit was used to convert the raw reads to a FASTQ format. *BBDuk* was used to find and remove adapter sequences from the 3’ end of reads, and rRNA reads were identified and removed using *BBMap* [97]. The cleaned Ribo-Seq reads were aligned to the reference genome by *HISAT2* [98]. The actively translating ORFs were detected and quantified by *Ribotricer*, which considers the periodicity of ORF profiles and provides multiple options for customization (we used the recommended parameters for yeast) [15]. The gene/ORF with mean counts across 302 Ribo-Seq samples higher than 0.3 was consider to have translation evidence.

### Visualization and gene function exploration

As proof-of-concept for the utility of these data, we used the MOG platform [51] to provide examples of co-expression and functional inference. We first created a MOG project that combined: 1) the levels of expression of each gene and ORF in the SGD+ORF dataset across 3,457 conditions, 2) gene and ORF metadata, and 3) sample metadata. The gene and ORF metadata includes: functional annotations (from SGD); MCL cluster memberships with GO enrichment analysis; mean expression levels for RNA-Seq and ribosomal profiling; ribosomal binding evidence; genome location relative to UTRs and CDSs; GC content; length; genomic positional coordinates, orientation; and phylostratal assignment. We then added metadata to the MOG project about each sample and study from NCBI-SRA, including: study ID, title, summary, reference, design description, library construction protocol, sequencing apparatus; sample title, experimental attributes, number of replicates; replicate name, sequencing depth, base coverage.

## Authors’ contributions

JL and EW conceived of the project and drafted the manuscript. All authors contributed to the manuscript. JL carried out the design of the study and performed the statistical analysis. US participated in the visualization on MOG and provided Ribo-Seq analysis code and guidance. ZA contributed to the phylostrata analysis. All authors read and approved the final manuscript.

## Acknowledgements

We thank Kevin Bassler and Pramesh Singh at University of Houston for insightful discussion about clustering analysis, and Manhoi Hur and Nishanth Sivakumar for developing the MCL Spark pipeline used in this study. We are grateful to Arun Seetharam and Priyanka Bhandary for helpful discussion on RNA-Seq alignment. This work was enabled by an award to ESW, AS and US from Extreme Science and Engineering Discovery Environment (XSEDE) supported by National Science Foundation ACI-1548562, and by Condo Cluster at Iowa State University.

## Supplementary material

Supplementary Materials.pdf: include all supplementary figures, tables and description.

All supplementary data (including MOG files, raw count data, cluster information, UTR results, phylostratr heatmap, Ribo-Seq metadata and results) are available at https://datahub.io/lijing28101/yeast_supplementary

MOG file of *S. cerevisiae* RNA-Seq expression (*S.cerevisiae_RNA-seq_3457_27.mog*): http://metnetweb.gdcb.iastate.edu/MetNet_MetaOmGraph.htm

MetaOmGraph software: https://github.com/urmi-21/MetaOmGraph

Data processing code: https://github.com/lijing28101/yeast_supplementary

