## Supplementary_Material.pdf for "Landscape of the Dark Transcriptome Revealed through Re-mining Massive RNA-Seq Data"

### Supplementary Materials

#### Section 1. Orphan gene identification

To stratify the phylostrata level for each gene/ORF, we used a total of 124 species, 17 of which are speices within *Saccharomycotina*. Specifically, we used 10 species with high quality genomes for the clade *Saccharomycotina*, and 7 species with high quality genomes for the clade *Saccharomyces ssp*. To account for potential incomplete gene annotation in *Saccharomyces,* we included as input data the translation products of all ORFs for each *Saccharomyces* species, as well as all the annotated proteins. Thus, if a (translated) ORF in *S. cerevisiae* had homology to a (translated) ORF, it was assigned to that clade, rather than as an orphan-ORF. This provided a stringent and more conservative determination of orphan-ORFs.

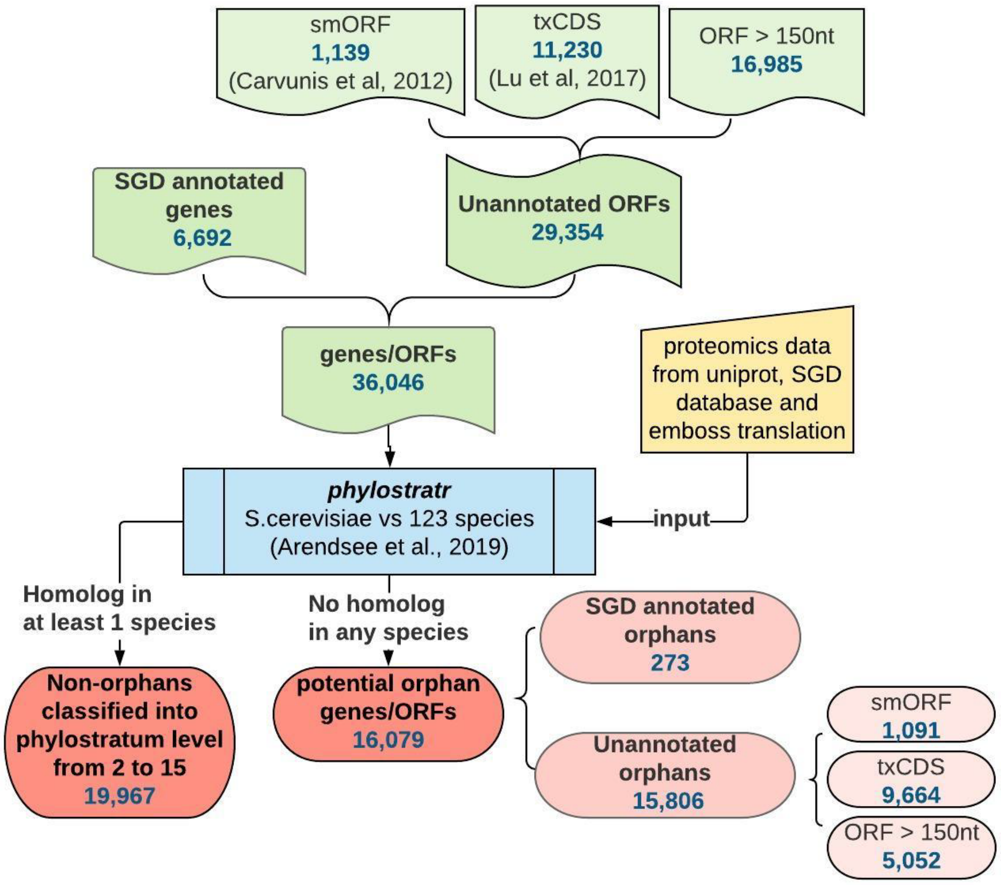

Figure S1. **Orphan-ORF detection by *phylostratr* (Arendsee et al, 2019).** The phylostrata of the 36,046 genes and ORFs in *S. cerevisiae* (determined by *phylostratr* (Arendsee et al., 2019)) by pairwise BLAST of the proteins of *S. cerevisiae* compared to other 123 species (see *species.xlsx*); E-value cutoff to infer homology, 0.001. The default NCBI clade tree was used except for species within *Saccharomyces*. For these species, we revised the clade tree manually to reflect current knowledge of yeast phylogeny (Borneman et al., 2015). Genes/ORFs were assigned to the phylostratum level associated with the deepest clade with inferred homolog. Genes/ORFs were inferred to be orphans if they had no homologs in the any of the other 123 species. Note that, for ease of comparison among the literature, throughout this study when we are specifically comparing groups of ORFS, we refer to "smORFs" (Carvunis et a., 2012) and "txCDS" (Lu et al.,2017) by the names designated in the papers that described them; the ORFs we identify in this study we designate as ORFS >150 nt. If we are not dividing into groups, we refer to them as ORFs. smORF, small ORFs with translation evidence identified by ribosomal profiling (Carvunis et al., 2012); ORFs, unannotated ORFs of ≥ 150 nt (excluding smORFs); txCDS, ORFs < 150 nt with transcription evidence identified by TIF-seq (Lu et al., 2017);

Section 2. CDS/ORF length and GC content across phylostrata

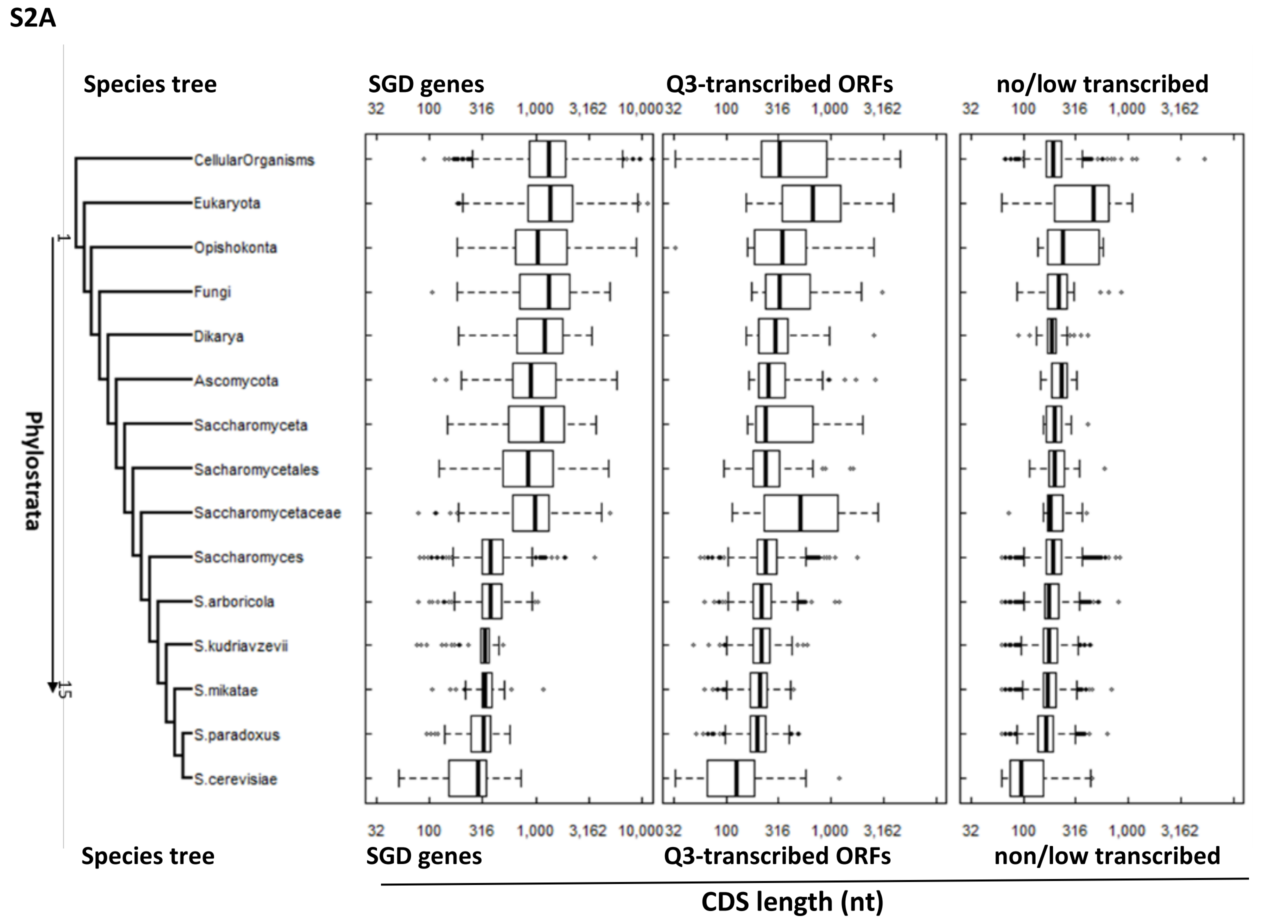

(Continues on following page)

**S2B.**

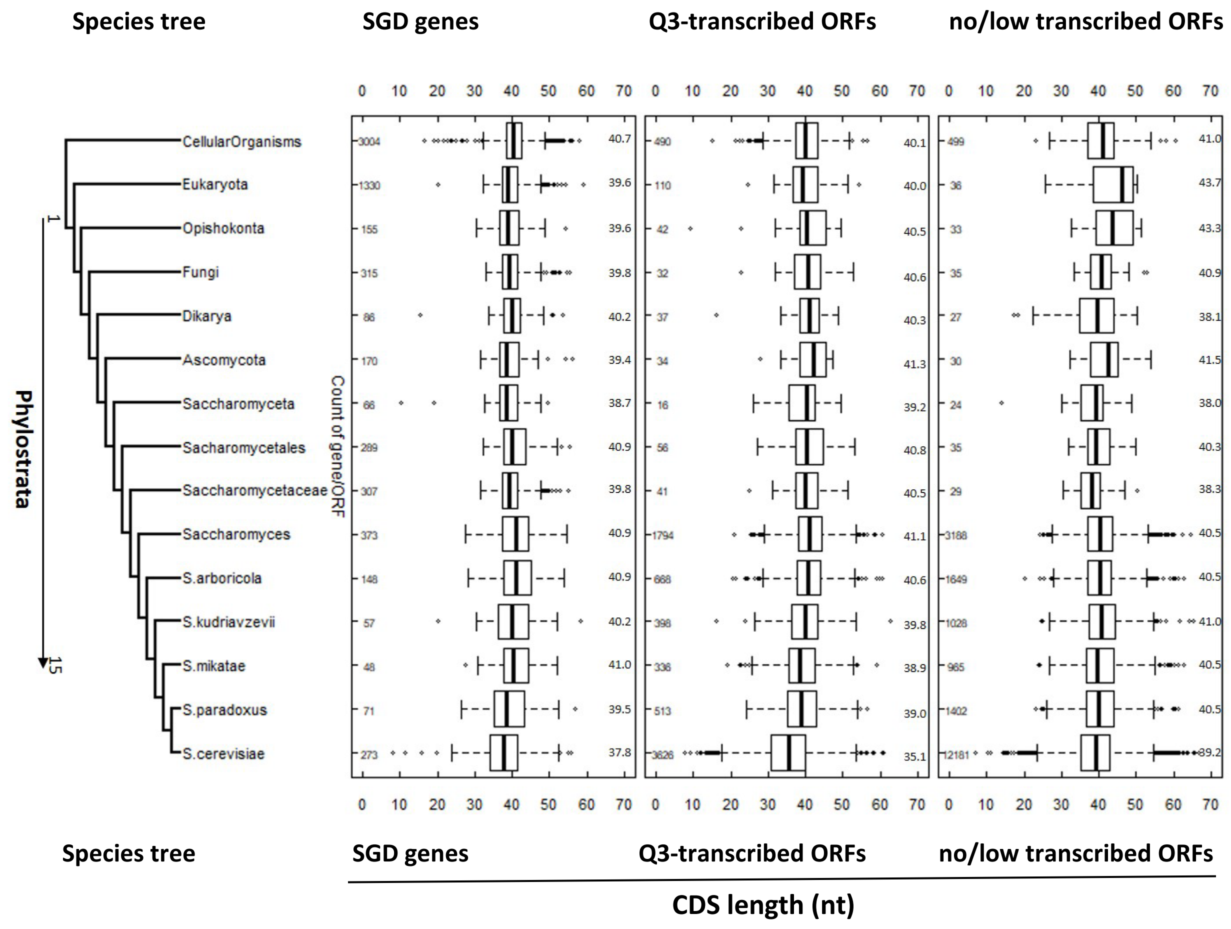

Figure S2. **CDS length and GC content of transcripts note fig 2B by phylostrata for SGD annotated genes, transcribed unannotated ORFs and non-transcribed/minimally transcribed unannotated ORFs.** From left, phylogenetic tree; first vertical panel, SGD-annotated genes (6,692); second vertical panel, Q3-transcribed ORFs, first quantile of most highly expressed unannotated ORFs (based on mean expression value/sample) (8,193); third vertical panel, no/low expressed unannotated ORFs (21,161). (A) Boxplots of CDS or ORF length. Orphan SGD genes and Q3-transcribed orphan-ORFs are significant shorter than conserved genes/ORFs. (B) Boxplots of GC content of CDS or ORF. No statistically significant for GC content between orphan and conserved genes/ORFs. The numbers of genes and ORFs in each phylostratum level are indicated in small font on left side of each vertical panel (numbers of genes and ORFs in each phylostratum level are identical for panel A and B).

Section 3. RNA-Seq expression for different categories of transcripts

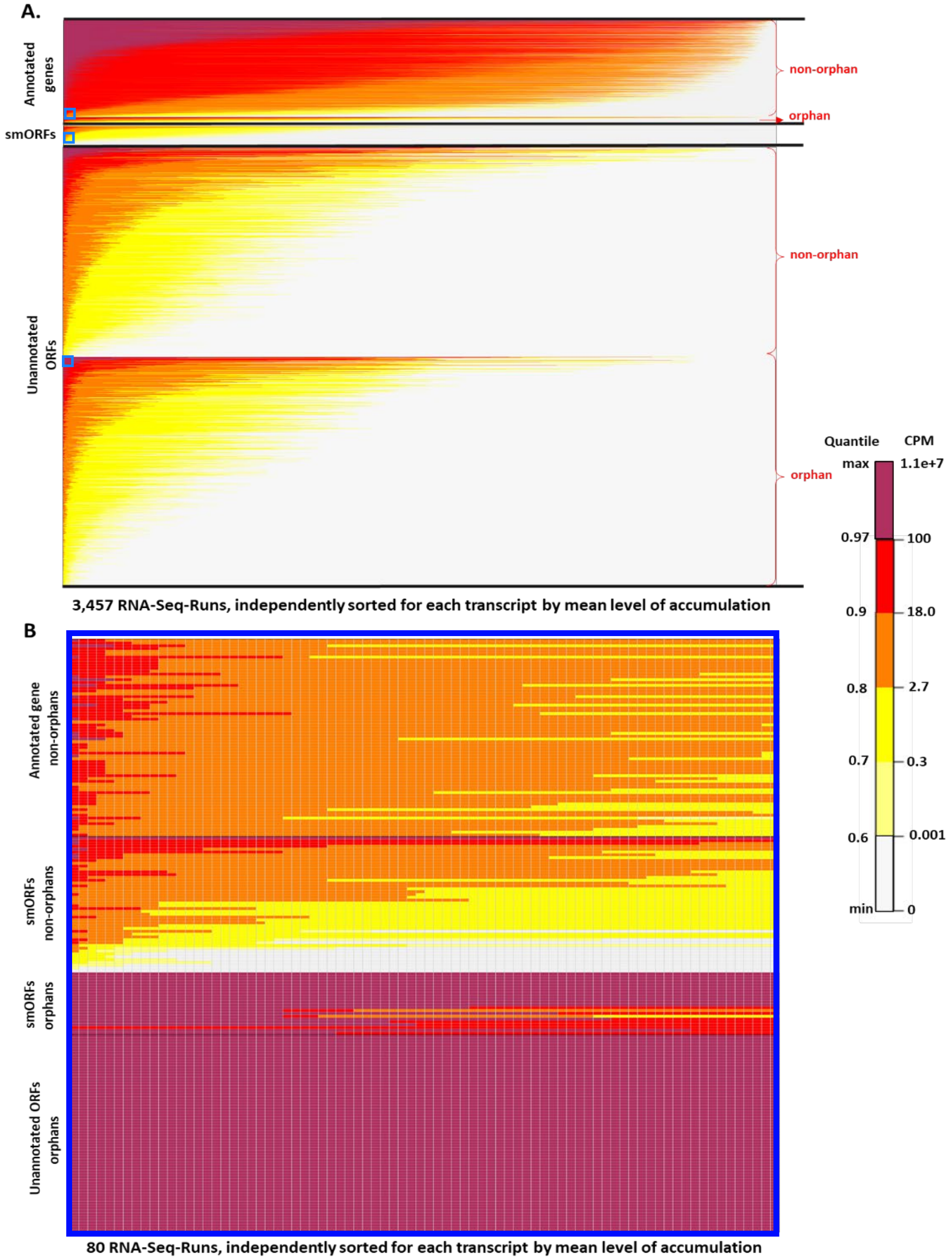

Figure S3. **RNA-Seq expression heatmap for all transcripts across 3,457 RNA-Seq runs.** Each group of transcripts is ordered independently by its mean cpm across 3,457 runs, in descending order. Each transcript is independently sorted (descending order) by cpm in sample. *EdgeR* was used for transcript normalization. (A) Heatmap of expression values for transcripts across 3,457 samples. Panels (top to bottom): 6,692 SGD-annotated genes: 6,419 non-orphan genes; 273 orphan genes; smORFs: 1,139 (Carvunis et al., 2012); all unannotated ORFs (28,215) (including 15,805 orphan-ORFs, 11,942 genus-specific ORFS, and 1,606 more highly conserved ORFS). Note: the proportion of orphans among the annotated orphan genes is very small and hard to distinguish. (B) High-resolution (blown-up) heatmap of the area *within the tiny blue boxes in panel A* (Areas marked in panel A are larger than actual area shown in panel B, for visibility). Each square corresponds to the expression value for a gene or ORF in a given sample. Panels (top to bottom): non-orphan SGD-annotated genes (#=70); non-orphan smORFs (#=48); orphan smORFs (#=22); orphan-ORFs (#=70).

To evaluate the distribution of mean expression values among the annotated genes and Q3-transcribed unannotated ORFs, we made density plots for mean expression across the 3,457 samples (Figure 4 in manuscript). We grouped SGD-annotated genes and ORFs according to their assignment to orphan versus all non-orphan ("conserved") phylostrata. The orphan-ORFs have two density peaks (Figure 4 in manuscript). When the ORFs are sub-divided further and then plotted, (Figure S4), it is clear that smORFs have lower expression other groups of ORFs. Because the smORFs are generally shorter, we evaluated whether the lower expression might be due to a bias of *kallisto* in aligning/quantifying shorter ORFs. To do this, we divided each type of ORF into four quantiles according to their length in nt, with Q1 being the shortest (Table S1). A density plot (Figure S5) shows that every length range of smORFs has a lower expression than any other group of ORFs; e.g., the mean of Q4 smORF expression is lower than any for txCDS despite that Q1 and Q2 txCDSs are shorter in length. Hence, the low expression of smORFs does not appear to be due to a technical bias in quantification of shorter reads.

A

| conserved SGD-annotated genes | | | orphan SGD-annotated genes | | all conserved-ORFs | | all orphan-ORFs | |
|---|---|---|---|---|---|---|---|---|
| quantile(%) | mean cpm | number of genes | quantile(%) | number of genes | quantile(%) | number of ORFs | quantile(%) | number of ORFs |
| min | 1.64E-02 | 6419 | 0.73 | 271 | 0.09 | 4564 | 7.64 | 3350 |
| 5 | 2.27E+00 | 6098 | 59.34 | 111 | 40.22 | 2731 | 58.96 | 1489 |
| 10 | 5.62E+00 | 5777 | 73.99 | 71 | 68.64 | 1433 | 81.52 | 671 |
| 25 | 1.40E+01 | 4814 | 85.71 | 39 | 83.32 | 763 | 92.06 | 289 |
| 50 | 2.71E+01 | 3210 | 90.84 | 25 | 90.10 | 453 | 95.81 | 153 |
| 75 | 5.73E+01 | 1605 | 95.97 | 11 | 94.24 | 264 | 98.01 | 73 |
| 90 | 1.31E+02 | 642 | 97.44 | 7 | 97.22 | 128 | 99.03 | 36 |
| 95 | 2.32E+02 | 321 | 99.27 | 2 | 98.55 | 67 | 99.45 | 21 |
| max | 3.13E+05 | 1 | NA | 0 | NA | 0 | NA | 0 |

B

| conserved-ORFs (≥150nt) | | orphan-ORFs (≥150nt) | | conserved-smORF | | orphan-smORF | | conserved-txCDS | | orphan-txCDS | |
|---|---|---|---|---|---|---|---|---|---|---|---|
| quantile(%) | number of ORFs | quantile(%) | number of ORFs | quantile(%) | number of ORFs | quantile(%) | number of ORFs | quantile(%) | number of ORFs | quantile(%) | number of ORFs |
| 0 | 4386 | 0 | 1625 | 8.33 | 45 | 25.39 | 815 | 0 | 133 | 0 | 910 |
| 39.47 | 2655 | 41.29 | 954 | 81.25 | 10 | 95.51 | 50 | 48.87 | 68 | 46.26 | 489 |
| 68.01 | 1403 | 71.32 | 466 | 91.67 | 5 | 98.08 | 22 | 77.44 | 30 | 79.56 | 186 |
| 82.95 | 748 | 86.65 | 217 | 93.75 | 4 | 99.18 | 10 | 90.23 | 13 | 93.08 | 63 |
| 89.88 | 444 | 92.55 | 121 | 97.92 | 2 | 99.45 | 7 | 94.74 | 7 | 97.25 | 25 |
| 94.05 | 261 | 96.25 | 61 | max | 1 | 99.82 | 3 | 98.50 | 2 | 99.01 | 9 |
| 97.15 | 125 | 98.22 | 29 | max | 1 | max | 1 | 98.50 | 2 | 99.34 | 6 |
| 98.50 | 66 | 98.89 | 18 | max | 1 | max | 1 | NA | 0 | 99.78 | 2 |
| max | 1 | NA | 0 | NA | 0 | NA | 0 | NA | 0 | NA | 0 |

Table S1. **Quantile of mean expression level for each group of transcripts.** Groups of transcripts are compared relative to the quantiles of mean expression of conserved SGD-annotated genes. Mean expression level is the mean across the 3,457 RNA-Seq samples (A) Conserved SGD-annotated genes, orphan SGD-annotated genes, conserved ORFs, orphan-ORFs. (B) Subdivisions of ORFS: conserved-ORFs (≥150nt), orphan-ORFs (≥150nt), conserved-smORF, orphan-smORF, conserved-txCDS, and orphan-txCDS.

| | | min-25% | 25%-50% | 50%-75% | 75%-max |
|---|---|---|---|---|---|
| smORFs | mean length (nt) | 36 | 45 | 58 | 107 |
| | number of ORFs | 269 | 253 | 313 | 304 |
| txCDSs | mean length (nt) | 70 | 88 | 110 | 135 |
| | number of ORFs | 283 | 252 | 262 | 246 |
| ORFs (≥150nt) | mean length (nt) | 168 | 203 | 250 | 634 |
| | number of ORFs | 1,574 | 1,478 | 1505 | 1454 |
| orphan-ORFs (≥150nt) | mean length (nt) | 161 | 184 | 213 | 274 |
| | number of ORFs | 451 | 366 | 417 | 391 |
| conserved-ORFs (≥150nt) | mean length (nt) | 173 | 215 | 266 | 747 |
| | number of ORFs | 1,162 | 1,070 | 1,068 | 1,086 |

Table S2. **Quantile of mean length for each group of transcripts.** smORFs, txCDSs, and ORFs (≥150nt) were divided into quantiles according to mean length of ORF. Mean length, mean length for each quantile range; number of ORFs, number of ORFs in each quantile range.

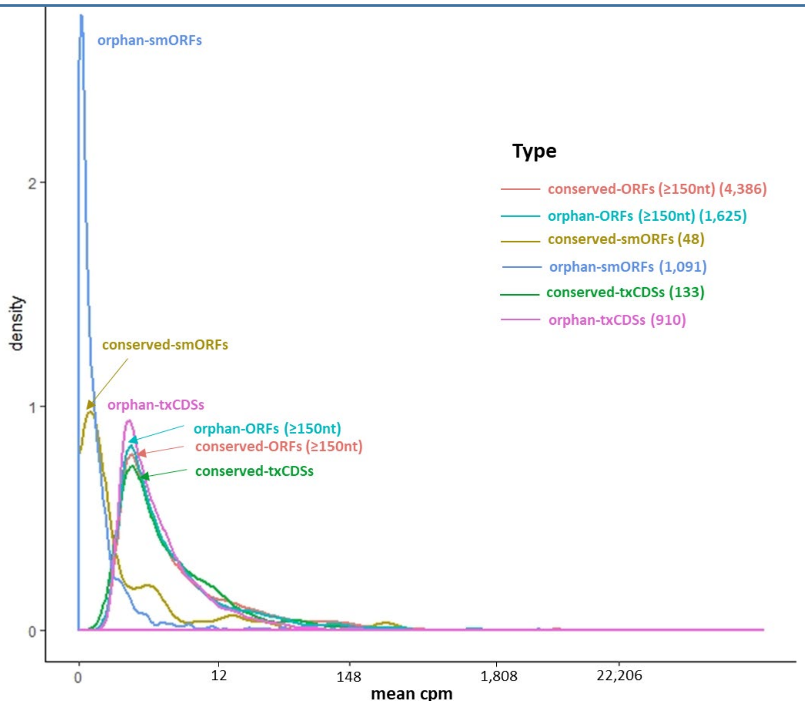

Figure S4. **Density plot for mean expression level of transcripts separated by different types of ORFs.** X-axis, *edgeR*-normalized mean expression of ORFs across 3,457 samples. The area under the curve of the density function represents the probability of a range of mean cpm.

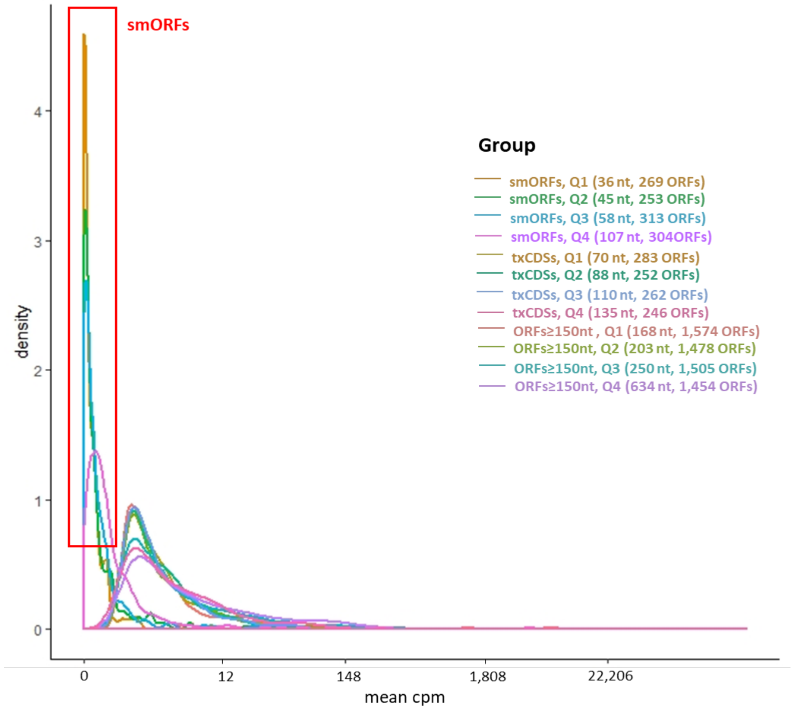

Figure S5. **Density plot showing mean expression level for each length range of different groups of ORFs.** X-axis, *edgeR*-normalized mean expression of transcripts in cpm across 3,457 samples. The area under the curve of density function represents the probability of a range of mean cpm. ORFs are classified by length: Q1 to Q4 refers to the range of length in nt, from short (Q1) to long (Q4). ORF mean length (nt) and number of ORFs in that category are given in parentheses.

Section 4. Transcription levels for ORFs relative to genomic context.

To investigate whether the trasncription level of unannotated ORFs was different relative to their genomic context, we divided the 29,354 ORFs into seven groups, according their relation to annotated CDSs. These groups are: 1) ORFs within the interval between annotated CDSs, 2) ORFs overlapping two (or more) annotated CDSs that express in the same and reverse orientation compared to the ORF, 3) ORFs overlapping annotated CDSs that express in the reverse orientation compared to the ORF, 4) ORFs overlap annotated CDSs that express in the same orientation as the ORFs, 5) ORFs within two (or more) annotated CDSs which express in the same and reverse orientations comparing to the ORFs, 6) ORFs within annotated CDSs that express in the reverse orientation comparing to the ORFs, 7) ORFs within annotated CDSs that express in the same orientation as the ORFs. The mean expression level and number of ORFs in each group are shown in Figure S6. ORFs overlapping annotated CDSs have a higher median of mean transcription (Wilcoxon rank-sum test, p-value $< 0.001$). Of the 6,742 ORFs with transcription and translation evidence according to the mRNA-Seq and Ribo-Seq analysis, 31% are in the interval between annotated CDS, and most of these are orphan-ORFs (Figure 5 in manuscript). In contrast, among the 289 orphan-ORFs which have mean expression higher than 25% of the conserved SGD-annotated genes, the proportion of orphan-ORFs overlapping CDSs is significantly higher than other ORFs (p-value < 0.001, Fisher's exact test) (Figure S7).

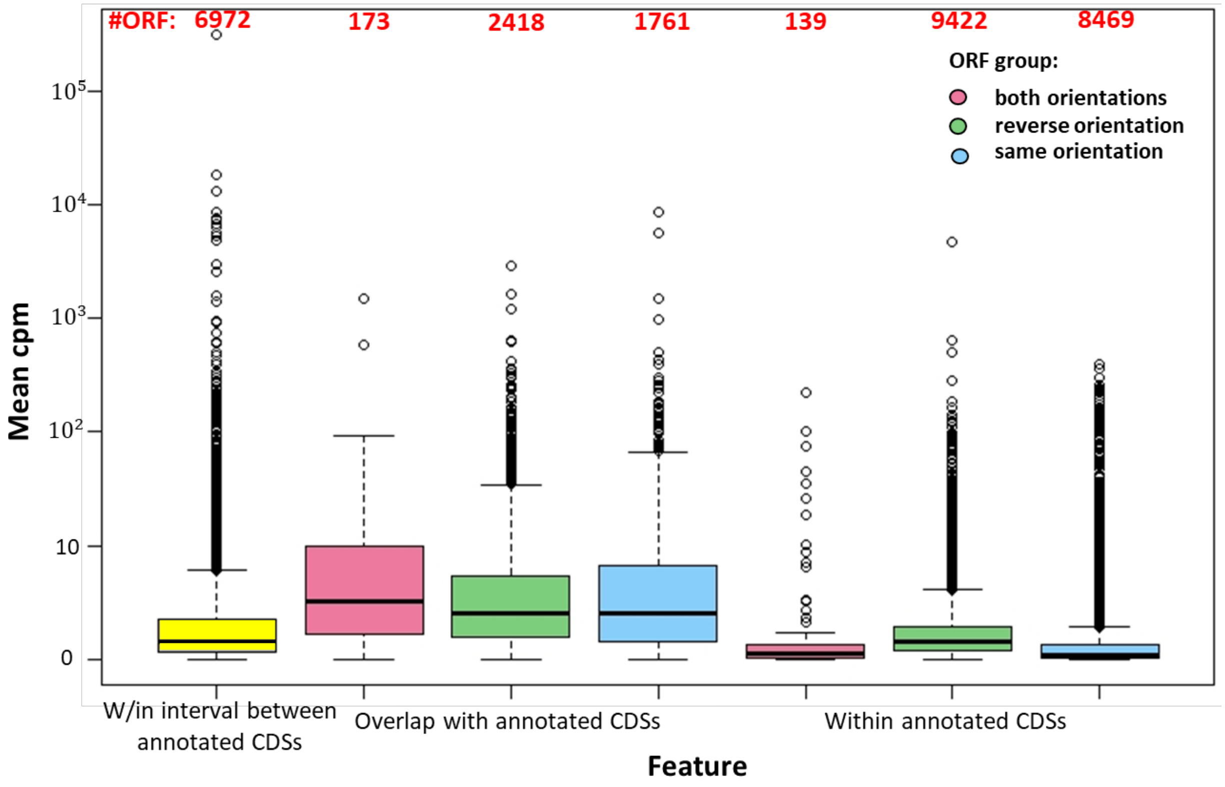

Figure S6. **Boxplot for mean expression level of ORFs, based on genomic context.** The median of mean expression is 5-fold greater for ORFs that overlap CDS. Most ORFs are located within annotated CDS. X-axis, classes of ORFs in relation to annotated CDSs. Y-axis, mean cpm across 3,457 RNA-Seq samples. Green boxes, overlaps or within annotated CDSs that express in the reverse (convergent or divergent) orientation. Blue boxes, ORFs overlap or within annotated CDSs that express in the same orientation (co-orientation). Pink boxes, ORFs overlap or are within two (or more) annotated CDSs that express in both the reverse and co-orientation. Yellow box, within the interval between two annotated CDSs. (See Figure S7 for orientation explanation)

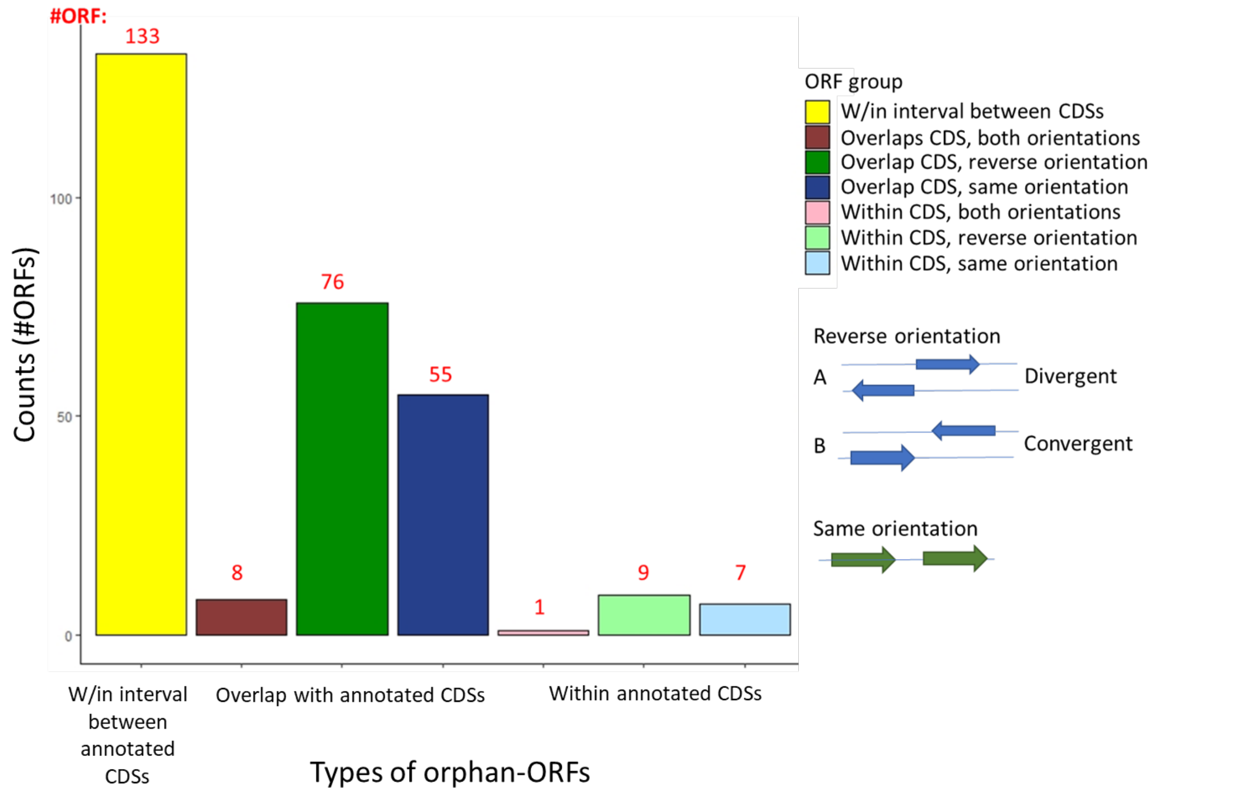

Figure S7. **Transcription evidence for orphan-ORFs with mean expression more than 25% of the conserved annotated genes, based on genomic context.** The 289 orphan-ORFs with mean expression higher than 25% of the conserved SGD-annotated genes were divided into different groups according to their relationship to adjacent or overlapping CDS (as in Figure S6). The number is shown for each group. Although in general most ORFs are located within CDS (Figure S6), the vast majority of these highly-expressed orphan-ORFs are either overlapping or outside annotated CDS. X-axis, groups of ORFs. Y-axis, counts of orphan-ORFs.

Section 5. Ribo-Seq analysis

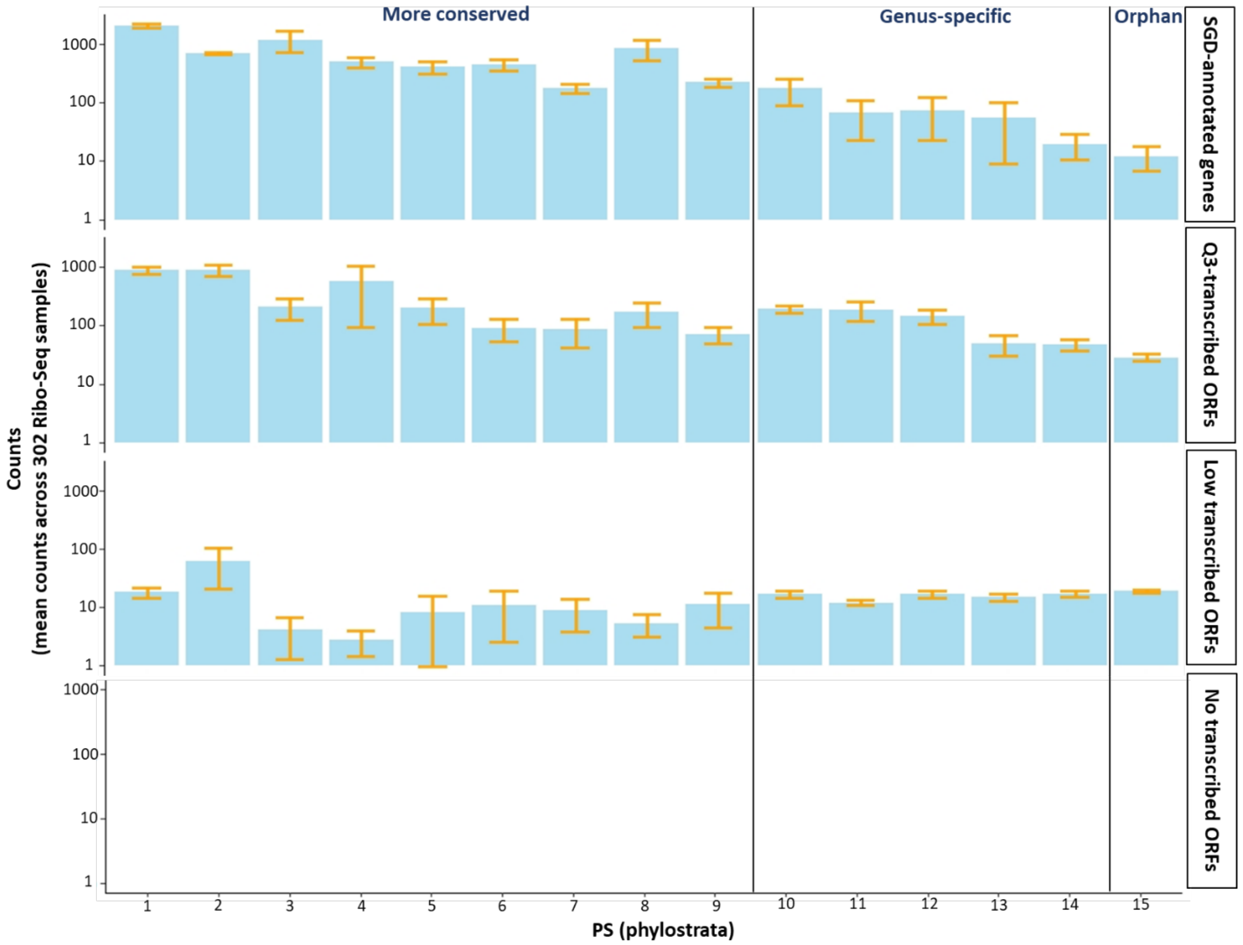

Figure S8. **Mean raw counts across 302 Ribo-Seq samples for different groups of genes/ORFs.** Translation evidence for the Q3 ORFs is similar in counts to that of the annotated genes. X-axis, PS (phylostrata) from 1 to 15 means from ancient to orphan. Orphan, PS=15; genus-specific, PS=10-14; more conserved, PS=1-9. Y-axis, mean raw counts across 302 Ribo-Seq samples for each group, the y-axis for no transcribed ORFs is different than other three groups. Yellow segment, standard error bar. An ORF is considered to have translation evidence if the mean value is greater than 0.3. Note: only 49 ORFs are non-transcribed, all in PS=15. Their mean count is less than 1, and the bar is too small to shown in the figure.

Section 6. Data, metadata download and RNA-Seq raw data processing.

The RNA-Seq data analysis workflow is shown as Figure S9. All the transcriptomic metadata and raw data were collected from public database. First, we collected SRR run IDs from (National Center for Biotechnology Information-Sequence Read Archive) (NCBI-SRA) using SRA advanced search builder. We chose all runs with *S.cerevisiae* taxon ID 4932, Illumina platform, and paired layout, and then filtered out the runs with miRNA-Seq, ncRNA-Seq, and RIP-Seq library strategy. In all, we collected 3,457 RNA-Seq runs from 177 studies. We used R packages *SRAdb* (Zhu et al., 213) and *GEOmetadb* (Zhu et al., 2008) to download RNA-Seq metadata from SRA and GEO database. *SRA toolkit* was used to download the raw RNA-Seq reads from the SRA database.

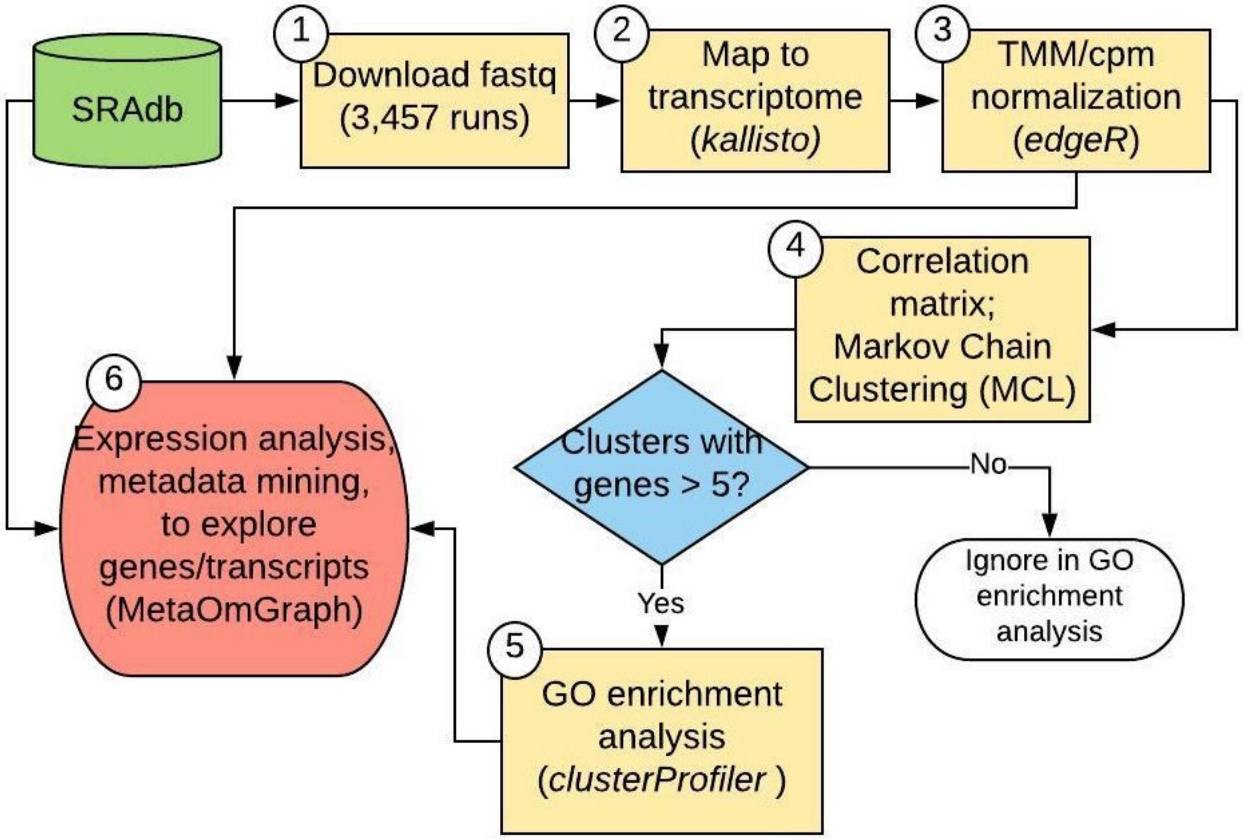

Figure S9. **RNA-Seq data analysis workflow.** 1) Raw reads and metadata for 3,457 samples (runs) of RNA-Seq were downloaded from SRA. 2) RNA-Seq reads were mapped to a transcriptome of genes and ORFs and abundance was quantified by kallisto. 3) TMM normalization (Robinson & Oshlack, 2010) by edgeR was applied to minimize technical variability of the data, with cpm as units. 4) A Pearson correlation matrix was calculated and MCL clustering analysis was performed using a home script (See code in GitHub. 5) GO enrichment analysis was conducted for clusters of five or more transcripts. 6) Full data and metadata were interactively statistically analysed and visualized in MOG (Singh et al., 2020) to explore and infer functions of selected transcripts.

#### Section 7. Co-expression cutoff determinations, clustering, and GO enrichment

To determine a cutoff range for Pearson Correlation Coefficient (PCC), we evaluated the largest connected component and the network density for multiple PCC cutoffs (Figure S10.A). The largest connected component decreased with an increasing PCC cutoff, a linear decrease in the largest connected component occurs between 0.6 and 0.8. Network density rapidly increases when the cutoff is below 0.7, indicating that the available nodes become more densely connected with the increase in the network density below this cutoff. Thus, this method indicates a PCC cutoff of between 0.6 and 0.7. We also applied *findThreshold* function in *coexnet* (R package) (Henao et al., 2017), this method determines the optimal cutoff at 0.74 (Figure S10.B). Therefore, we tested the performance of 0.6, 0.7, and 0.8 as PCC cutoffs for the networks used in MCL clustering.

PCC co-expression matrices for the SGD and SGD+ORF datasets were transformed into a binary matrix by replacing the values of all correlations larger than the selected cutoff value by 1, and assigning the others as 0. The matrices were provided to MCL clustering software. A Spark JAVA script was used for network inference and MCL clustering (https://github.com/lijing28101/SPARK_MCL); this script is faster than R and can handle larger datasets. The power and inflation parameters for MCL analysis were set as 2.

For GO enrichment analysis of the resultant clusters, yeast genes were mapped to GO terms downloaded from the SGD database. The over-representation values for GO terms in each cluster were obtained by *enricher* in *clusterProfiler* (R package) (Yu et al., 2012); the maximal size of genes for a GO term was set as 500, and the minimal size was set as 1.

In a random test distribution of GO enrichment analysis (Mentzen and Wurtele, 2008), the experimental data has smaller p-values than any random data for networks derived using any of three cutoffs. However, cutoffs of 0.6 and 0.7 perform better, since the distance of experimental data from the random distribution is larger than for cutoff 0.8 (Figure S11). Then, we compared MCL analysis of networks based on cutoffs 0.6 and 0.7 using mean of the lowest p-value for each cluster of the experimental data (Mentzen and Wurtele, 2008), and the Z-score between the experimental data and random distribution data (Table S3). The mean of the lowest p- values for experimental data are very similar for these two cutoffs, but the Z-scores for GO molecular function (MF) and biological process (BP) of PCC 0.6 are better than those of 0.7. Moreover, PCC 0.6 cutoff resulted in more orphan genes in the clusters. Thus, we choose 0.6 as the MCL clustering cutoff for the further analysis.

A.

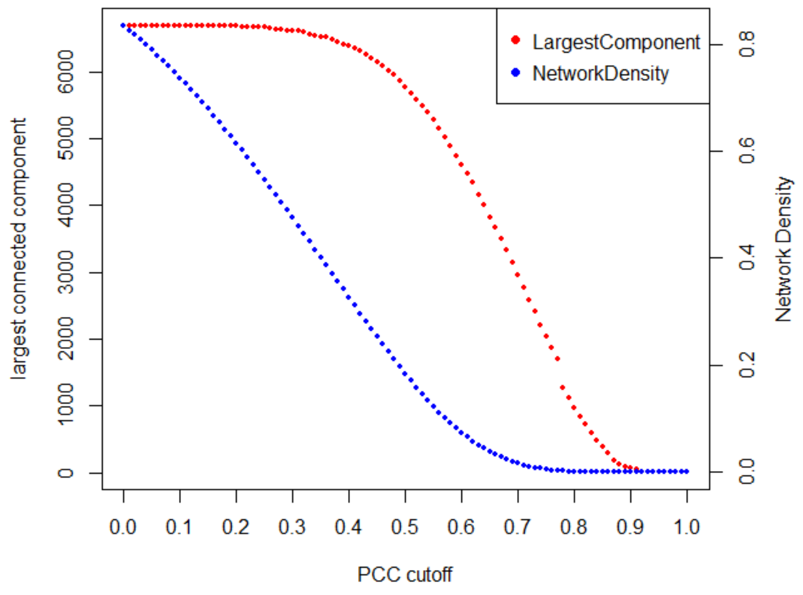

B.

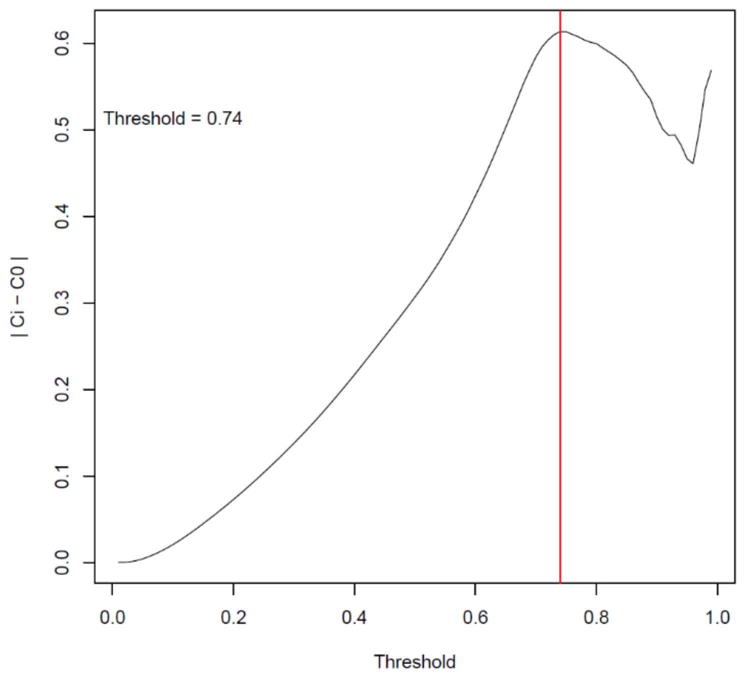

Figure S10. **Pearson correlation coefficient cutoff determination.** (A) Graph of the number of nodes in the largest connected component and network density for each Pearson correlation coefficient (PCC) cutoff. (B) The Pearson correlation coefficient cutoff determined by *coexnet* (Henao et al., 2017). X-axis, Pearson correlation coefficient (the optimized threshold is 0.74). Y-axis, the difference of clustering coefficient between the current threshold value under test and simulated random network.

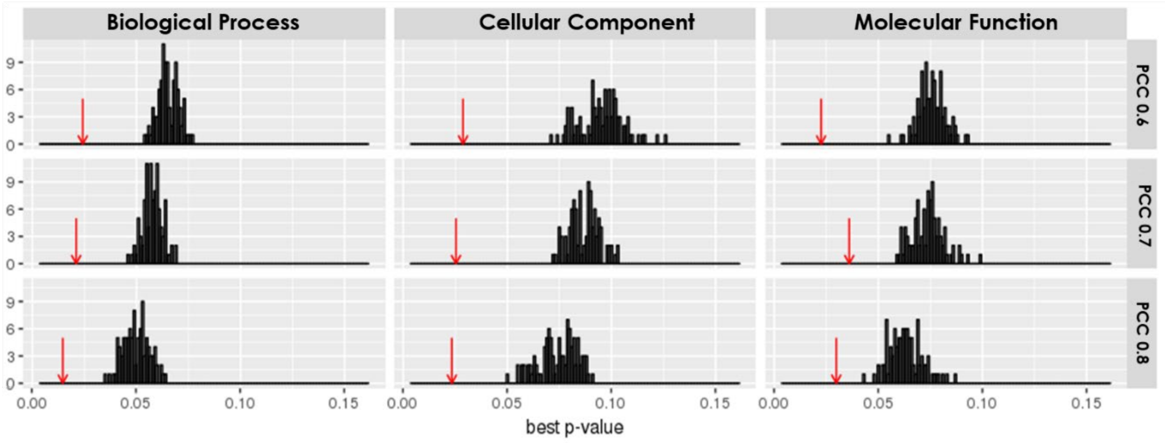

Figure S11. **Experimental versus random set distribution of GO enrichment analysis for SGD genes at three PCC cutoffs.** Red arrow, best p-value for the experimental data. Random distribution (black bars), results from 100 randomly-obtained datasets with the same size and number of clusters as the experimental data. The best p-value is the mean of best adjusted p-value of GO terms across all clusters in a random set or experiment data.

| Parameter | Ontology | PCC 0.6 | PCC 0.7 | PCC 0.8 |
|---|---|---|---|---|
| Mean of best p-value for experimental data | BP | 0.024 | 0.021 | 0.015 |
| | CC | 0.029 | 0.025 | 0.023 |
| | MF | 0.022 | 0.036 | 0.030 |
| Z-score* for experimental data vs random data | BP | 8.31 | 7.85 | 5.70 |
| | CC | 6.43 | 8.76 | 5.56 |
| | MF | 8.16 | 4.96 | 4.24 |
| Distance** for experimental data vs random data | BP | 0.041 | 0.036 | 0.035 |
| | CC | 0.068 | 0.063 | 0.052 |
| | MF | 0.053 | 0.038 | 0.033 |

Table S3. **Network PCC cutoff comparison for GO enrichment analysis.** BP, biological process; CC, cellular component; MF, molecular function.

*Z-score= $\frac{|\bar{p}_{expmin} - mean(\bar{p}_{ranmin})|}{sd(\bar{p}_{ran}min)}$

**distance= $|\bar{p}_{expmin} - median(\bar{p}_{ranmin})|$

#### Section 8. Comparison of normalization by *EdgeR* and *SCnorm*

Performance of the *EdgeR* and *SCnorm* normalization methods was assessed. *EdgeR* (Robinson et al., 2010) is a commonly applied package based on TMM normalization. We used M and A values to estimate scaling factors (Robinson et al., 2010), and then used counts per million (cpm) as units. *SCnorm* (Bacher et al., 2017) is a method developed for normalization of single cell RNA-Seq data. Groups are formed based on each gene's median expression; a quantile regression was applied to estimate scaling factors, based on each gene's count-depth relationship within each group; we are not aware of *SCnorm* having been applied to "standard' RNA-Seq data.

We evaluated the *edgeR* and *SCnorm* normalization by comparing boxplot of each samples among raw counts and two normalization methods (Figure S12), creating Pearson correlation matrices from the normalized data, applying a PCC cutoff of 0.6, followed by MCL clustering and GO term enrichment analysis. After MCL analysis, *SCnorm* normalization yielded more clusters and contained more transcripts, especially more orphan transcripts in the clusters than *edgeR* (Table S4). On the downside, *SCnorm* result had more transcripts in the largest cluster, and also included more small clusters with 5-10 genes and no GO term overrepresentation assignments.

We used ARI and Jaccard indices to compare the MCL clusters obtained from raw cpm data and after *edgeR* and *SCnorm* normalization. There was only partial overlap among the clusters obtained by these two methods. Clusters from the SGD+ORF dataset were less similar than SGD dataset (Table S5). GO enrichment analysis results for the two methods indicate more significant GO terms are found after *edgeR* normalization (Table S4). *edgeR* normalization performed better than *SCnorm* based on GO enrichment analysis of MCL clusters using datasets composed of only SGD annotated genes (SGD dataset) and composed of SGD-annotated genes and all unannotated ORFs (SGD+ORF dataset) (Figure S13). Hence, for our data, *edgeR* normalization yielded the best results, and was used for subsequent analysis.

| Gene list | Data | Total genes | Genes in clusters (total) | Clusters | Genes in largest cluster | Clusters with gene>5 | Total orphan | Orphans in clusters (total) | Orphans in cluster with gene>5 | Significant GO terms |
|---|---|---|---|---|---|---|---|---|---|---|
| SGD genes | Raw | 6,692 | 6,455 | 49 | 3,564 | 15 | 384 | 317 | 297 | |
| | edgeR | 6,692 | 4,351 | 226 | 2,509 | 49 | 384 | 155 | 125 | 3,160 |
| | scnorm | 6,692 | 4,622 | 289 | 2,573 | 67 | 384 | 242 | 170 | 2,154 |
| Combined transcripts | Raw | 14,885 | 13,639 | 194 | 10,884 | 44 | 4,010 | 3,435 | 3,314 | |
| | edgeR | 14,885 | 9705 | 544 | 3,328 | 89 | 4,010 | 2,647 | 2,336 | 2,205 |
| | scnorm | 14,885 | 10,630 | 562 | 4,658 | 133 | 4,010 | 3,012 | 2,752 | 1,950 |

Table S4. **Comparison of MCL clustering results for *edgeR* and *SCnorm* normalization.** Purple font, MCL results for SGD-annotated genes. Blue font, MCL results for SGD+ORF transcripts.

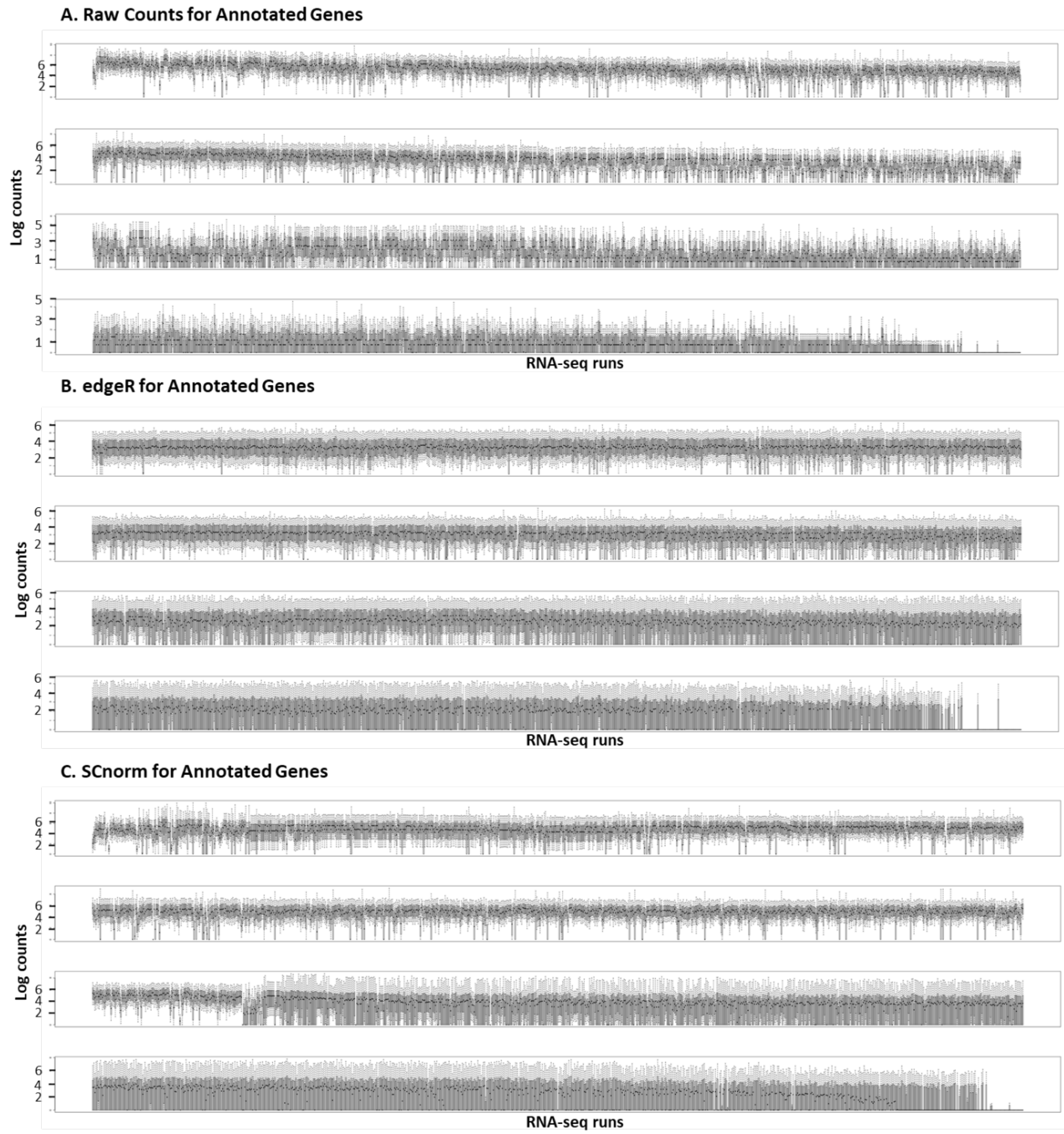

Figure S12. **Comparison of boxplots of each RNA-Seq run after *edgeR* and *SCnorm* normalization.** Values for SDG-annotated genes plotted across 3,457 samples are shown. X-axis, 3,457 RNA-Seq runs (split into 4 plots for ease of viewing). Y-axis, log transformed normalized counts. (A) Raw counts. (B) *edgeR*-normalized counts. (C) *SCnorm*-normalized counts.

| Similarity index | Raw vs TMM | Raw vs SCnorm | TMM vs SCnorm |
|---|---|---|---|
| | | **SGD** | |
| **ARI** | 0.140 | 0.144 | 0.586 |
| **Jaccard index** | 0.249 | 0.243 | 0.529 |
| | | **Combined Transcripts** | |
| **ARI** | 0.042 | 0.087 | 0.354 |
| **Jaccard index** | 0.181 | 0.212 | 0.313 |

Table S5. **Comparison of MCL clustering results for *edgeR* and *SCnorm* normalization using ARI and Jaccard indices**. These analyses evaluate similarity of clustering results across normalization methods. Adjusted index (ARI) (Rand et al., 1971); Jaccard index (Jaccard et al., 1901). SGD, SGD-annotated genes; Combined transcripts, SGD-annotated genes plus all ORFS (smORFs, txCDSs, + ORFs>150 nt). Raw, raw counts dataset; TMM, *EdgeR* normalization dataset; *SCnorm*, *SCnorm* normalization dataset

$$\overbrace{\widehat{ARI}}^{\text{Adjusted Index}} = \frac{\overbrace{\sum_{ij}\binom{n_{ij}}{2}}^{\text{Index}} - \overbrace{[\sum_i \binom{a_i}{2}\sum_j \binom{b_j}{2}]/\binom{n}{2}}^{\text{Expected Index}}}{\underbrace{\frac{1}{2}[\sum_i \binom{a_i}{2} + \sum_j \binom{b_j}{2}]}_{\text{Max Index}} - \underbrace{[\sum_i \binom{a_i}{2}\sum_j \binom{b_j}{2}]/\binom{n}{2}}_{\text{Expected Index}}}$$

where $n_{ij}, a_i, b_j$ are values from the contingency table.

$$\text{Jaccard index} = \frac{|A \cap B|}{|A \cup B|} = \frac{|A \cap B|}{|A| + |B| - |A \cap B|}.$$

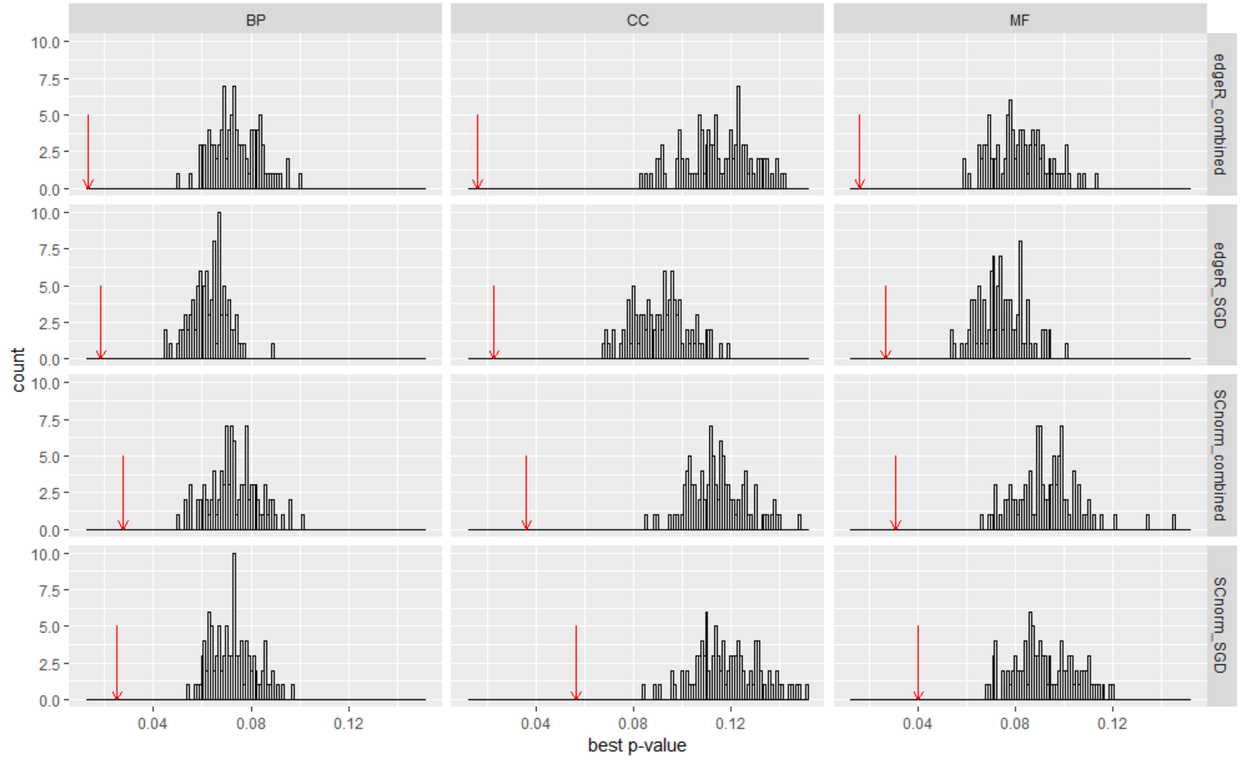

Figure S13. **Random set distribution of GO enrichment analysis for SGD genes and SGD+ORF combined transcripts for two normalization methods.** The red arrow is the best p-value for the experimental data. Random distribution, black bars, best p-value of GO terms from 100 randomly-obtained sets with the same size and number of clusters as the experimental data. Best p-value, mean of lowest adjusted p-value of GO terms across all clusters in a random set or in the experiment data (Mentzen and Wurtele, 2008).

#### Section 9. Study case: Cluster317

Cluster317 contains 25 highly conserved (phylostratum level 1 or 2) SGD-annotated genes and 5 *Saccharomyces*-specific ORFs. Twenty-three of the 25 conserved genes encode ribosomal proteins (P1 Alpha, large, or small subunits). Another gene encodes a zuotin, a ribosome-associated chaperone that functions in ribosome biogenesis. The remaining gene encodes guanylate kinase; a gene with nine edges in common with the other genes in this cluster of the co-expression network. In red alga, guanylate kinase and chloroplastic ribosomal proteins are co-regulated by the regulatory kinase, TOR, although the reason for this is not understood (Imamura et al., 2018). Each of the five ORFs share many highly correlated edges with other transcripts (Figure S14, left). GO enrichment analysis identified nine highly over-represented GO terms (Figure S14, right) each relating to ribosomes or translation. The expression pattern of the genes in Cluster317 is similar across the 3,457 RNA-Seq runs (Figure S14). The top panel of Figure S15 shows the expression level of Cluster317 in study ERP008497, which compares wild type and temperature-sensitive ubc9-1 mutant yeast in a DMSO control group vs. rapamycin treatment (Chymkowitch et al., 2015). In yeast, TOR, regulates ribosome biogenesis , along with cell proliferation, mRNA translation, responses to nutrients, autophagy, and mating (Cardenas et al., 1999). Thus, rapamycin represses the expression of yeast ribosomal protein genes. All the transcripts in this cluster have decreased expression following rapamycin treatment, irrespective of the yeast genotype. Combining this information, the five ORFs in Cluster317 might encode proteins associated with the ribosome, or be involved in the translation process.

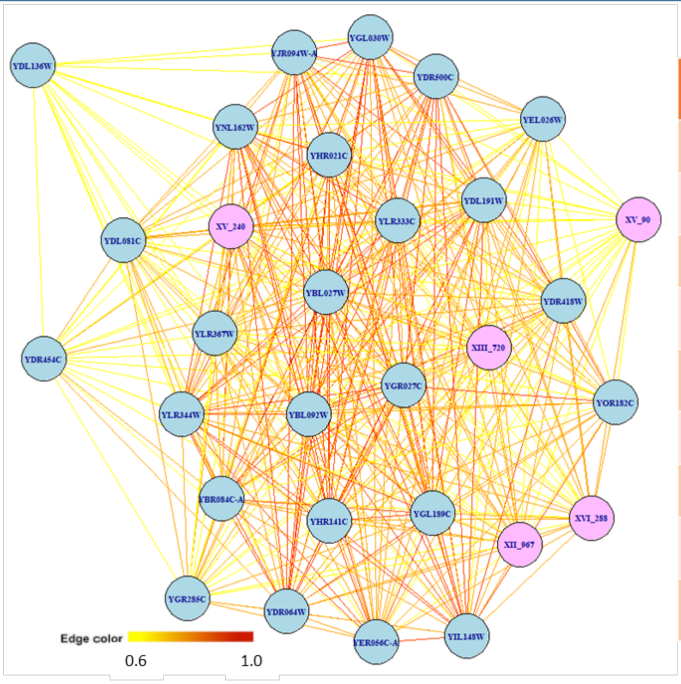

| Ontology | GO name | Adjust p-value |
|---|---|---|
| BP | cytoplasmic translation | 3.49E-32 |
| MF | structural constituent of ribosome | 4.73E-29 |
| CC | ribosome | 2.17E-28 |
| CC | intracellular ribonucleoprotein complex | 2.17E-28 |
| CC | cytosolic large ribosomal subunit | 1.32E-23 |
| BP | translation | 1.53E-22 |
| CC | intracellular | 3.03E-20 |
| CC | cytosolic small ribosomal subunit | 1.33E-08 |
| CC | preribosome, large subunit precursor | 7.12E-08 |

Figure S14. **Co-expression network and significant GO terms for Cluster317.** Cluster317 is composed of ribosomal related genes and five ORFs. Left, the network of Cluster317. Edge color, yellow to red, maps to Pearson correlations of 0.6 to 1. Light blue nodes, SGD-annotated genes; pink nodes, unannotated ORFs. Right, significantly enriched GO terms in Cluster317. Twenty-four of the 25 SGD-annotated genes in this cluster are assigned to GO terms in this table.

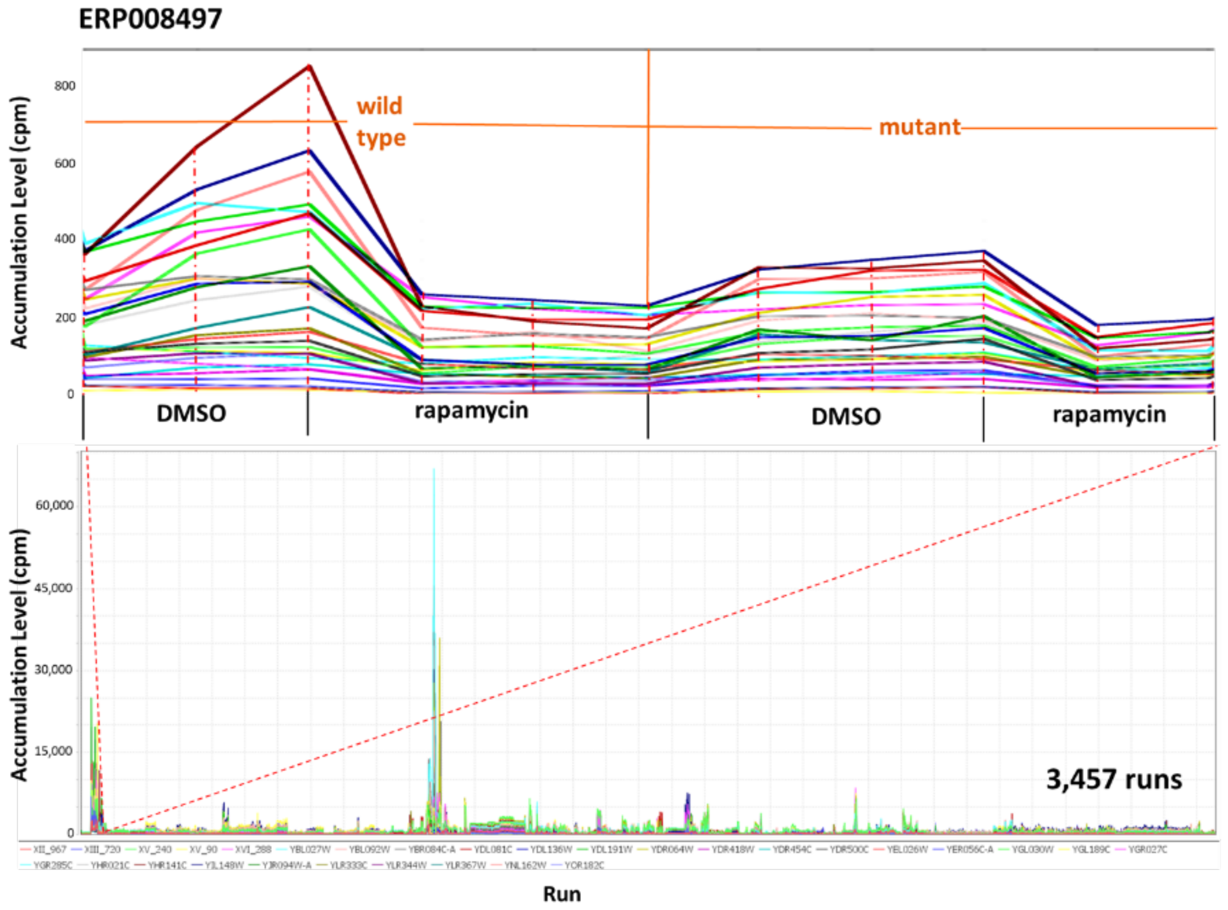

Figure S15. **The 41 genes and ORFs in Cluster317 are repressed in response to rapamycin.** Bottom panel, 3,457-samples. Expression pattern are similar for all transcripts. Top panel, high resolution of Study ERP008497: "Sumoylation of Rap1 mediates recruitment of the basal transcription machinery to promote transcription of Ribosomal Protein genes". Dimethyl sulfoxide (DMSO), control treatment. Rapamycin could repress expression of ribosomal protein.

#### Section 10. Other supplementary materials available online:

All of the supplementary data (include mog files, cluster information, UTR results, phylostratr heatmap, Ribo-Seq metadata and results): https://datahub.io/lijing28101/yeast_supplementary

MOG file of *Sacchromyces cerevisea* RNA-Seq expression (*S.cerevisiae_RNA-seq_3457_27.mog*): http://metnetweb.gdcb.iastate.edu/MetNet_MetaOmGraph.htm

MetaOmGraph software: https://github.com/urmi-21/MetaOmGraph

Data processing code: https://github.com/lijing28101/yeast_supplementary

Section 12. Saccharomyces sequence source

| file | download date | release date | version |
|---|---|---|---|
| *Saccharomyces cerevisiae* R64 gene model GFF3 | 10/17/2017 | 7/22/2017 | R64-1-1 |
| *Saccharomyces cerevisiae* R64 genomic sequence | 10/17/2017 | 7/22/2017 | R64-1-1 |
| *Saccharomyces cerevisiae* amino acid | 10/19/2017 | 7/22/2017 | R64-1-1 |
| smORF annotation | 10/19/2017 | 7/19/2012 | R56 |
| *Saccharomyces cerevisiae* R56 genomic sequence | 10/19/2017 | 4/6/2007 | R56 |
| *Saccharomyces arboricola* genomic sequence | 10/19/2017 | 1/15/2015 | SacArb1.0 |
| *Saccharomyces arboricola* protein sequence | 10/19/2017 | 1/15/2015 | SacArb1.0 |
| *Saccharomyces bayanus* genomic sequence | 10/19/2017 | 6/15/2016 | ASM16703v1 |
| *Saccharomyces bayanus* protein sequence | | | |
| *Saccharomyces eubayanus* genomic sequence | 10/19/2017 | 7/30/2017 | SEUB3.0 |
| *Saccharomyces eubayanus* protein sequence | 10/19/2017 | 7/30/2017 | SEUB3.0 |
| *Saccharomyces kudriavzevii* genomic sequence | 10/19/2017 | 8/4/2014 | IFO1802_v1.0 |
| *Saccharomyces kudriavzevii* protein sequence | 10/19/2017 | 6/15/2016 | IFO1802_v1.0 |
| *Saccharomyces mikatae* genomic sequence | 10/19/2017 | 8/1/2014 | ASM16697v1 |
| *Saccharomyces mikatae* protein sequence | | | |
| *Saccharomyces paradoxus* genomic sequence | 10/19/2017 | 6/15/2016 | ASM16695v1 |
| *Saccharomyces paradoxus* protein sequence | | | |
| *Saccharomyces uvarum* genomic sequence | 10/19/2017 | 6/15/2016 | ASM16699v1 |
| *Saccharomyces uvarum* protein sequence | | | |

| file | source |
|---|---|
| *Saccharomyces cerevisiae* R64 gene model GFF3 | ftp://ftp.ensembl.org/pub/release-90/gff3/saccharomyces_cerevisiae/Saccharomyces_cerevisiae.R64-1-1.90.gff3.gz |
| *Saccharomyces cerevisiae* R64 genomic sequence | ftp://ftp.ensembl.org/pub/release-90/fasta/saccharomyces_cerevisiae/dna/Saccharomyces_cerevisiae.R64-1-1.dna_sm.toplevel.fa.gz |
| *Saccharomyces cerevisiae* amino acid | ftp://ftp.ensembl.org/pub/release-90/fasta/saccharomyces_cerevisiae/pep/Saccharomyces_cerevisiae.R64-1-1.pep.all.fa.gz |
| smORF annotation | Carvunis at al., 2012 |

| | |
|---|---|
| *Saccharomyces cerevisiae* R56 genomic sequence | https://downloads.yeastgenome.org/sequence/S288C_reference/genome_releases/S288C_reference_genome_R56-1-1_20070406.tgz |
| *Saccharomyces arboricola* genomic sequence | ftp://ftp.ncbi.nlm.nih.gov/genomes/all/GCF/000/292/725/GCF_000292725.1_SacArb1.0/GCF_000292725.1_SacArb1.0_genomic.fna.gz |
| *Saccharomyces arboricola* protein sequence | ftp://ftp.ncbi.nlm.nih.gov/genomes/all/GCF/000/292/725/GCF_000292725.1_SacArb1.0/GCF_000292725.1_SacArb1.0_protein.faa.gz |
| *Saccharomyces bayanus* genomic sequence | ftp://ftp.ncbi.nih.gov/genomes/genbank/fungi/Saccharomyces_bayanus/latest_assembly_versions/GCA_000167035.1_ASM16703v1/GCA_000167035.1_ASM16703v1_genomic.fna.gz |
| *Saccharomyces bayanus* protein sequence | Annotated and created by Braker |
| *Saccharomyces eubayanus* genomic sequence | ftp://ftp.ncbi.nih.gov/genomes/genbank/fungi/Saccharomyces_eubayanus/latest_assembly_versions/GCA_001298625.1_SEUB3.0/GCA_001298625.1_SEUB3.0_genomic.fna.gz |
| *Saccharomyces eubayanus* protein sequence | ftp://ftp.ncbi.nih.gov/genomes/genbank/fungi/Saccharomyces_eubayanus/latest_assembly_versions/GCA_001298625.1_SEUB3.0/GCA_001298625.1_SEUB3.0_protein.faa.gz |
| *Saccharomyces kudriavzevii* genomic sequence | ftp://ftp.ncbi.nih.gov/genomes/genbank/fungi/Saccharomyces_kudriavzevii/latest_assembly_versions/GCA_000167075.2_Saccharomyces_kudriavzevii_strain_IFO1802_v1.0/GCA_000167075.2_Saccharomyces_kudriavzevii_strain_IFO1802_v1.0_genomic.fna.gz |
| *Saccharomyces kudriavzevii* protein sequence | ftp://ftp.ncbi.nih.gov/genomes/genbank/fungi/Saccharomyces_kudriavzevii/latest_assembly_versions/GCA_000167075.2_Saccharomyces_kudriavzevii_strain_IFO1802_v1.0/GCA_000167075.2_Saccharomyces_kudriavzevii_strain_IFO1802_v1.0_protein.faa.gz |
| *Saccharomyces mikatae* genomic sequence | ftp://ftp.ncbi.nih.gov/genomes/genbank/fungi/Saccharomyces_mikatae/latest_assembly_versions/GCA_000166975.1_ASM16697v1/GCA_000166975.1_ASM16697v1_genomic.fna.gz |
| *Saccharomyces mikatae* protein sequence | Annotated and created by Braker |
| *Saccharomyces paradoxus* genomic sequence | ftp://ftp.ncbi.nih.gov/genomes/genbank/fungi/Saccharomyces_paradoxus/latest_assembly_versions/GCA_000166955.1_ASM16695v1/GCA_000166955.1_ASM16695v1_genomic.fna.gz |
| *Saccharomyces paradoxus* protein sequence | Annotated and created by Braker |

| | |
|---|---|
| *Saccharomyces uvarum* genomic sequence | ftp://ftp.ncbi.nih.gov/genomes/genbank/fungi/Saccharomyces_uvarum/latest_assembly_versions/GCA_000166995.1_ASM16699v1/GCA_000166995.1_ASM16699v1_genomic.fna.gz |
| *Saccharomyces uvarum* protein sequence | Annotated and created by Braker |

*Section 13.* 124 species for *phylostratr*

| Species | PS | Phylostratum name |
|---|---|---|
| Aliifodinibius roseus | 1 | cellular organisms |
| Caldithrix sp. RBG_13_44_9 | 1 | cellular organisms |
| Aminobacterium colombiense DSM 12261 | 1 | cellular organisms |
| Treponema brennaborense DSM 12168 | 1 | cellular organisms |
| Thermodesulfatator indicus DSM 15286 | 1 | cellular organisms |
| Desulfurispirillum indicum S5 | 1 | cellular organisms |
| Denitrovibrio acetiphilus DSM 12809 | 1 | cellular organisms |
| Mesotoga prima | 1 | cellular organisms |
| Thermocrinis albus DSM 14484 | 1 | cellular organisms |
| Elusimicrobium minutum Pei191 | 1 | cellular organisms |
| Dictyoglomus turgidum DSM 6724 | 1 | cellular organisms |
| Caldisericum exile AZM16c01 | 1 | cellular organisms |
| Chloracidobacterium thermophilum B | 1 | cellular organisms |
| Candidatus Solibacter usitatus Ellin6076 | 1 | cellular organisms |
| Nitrospira japonica | 1 | cellular organisms |
| Fusobacterium mortiferum ATCC 9817 | 1 | cellular organisms |
| Hydrogenophilaceae bacterium CG1_02_62_390 | 1 | cellular organisms |
| Acidithiobacillales bacterium SM1_46 | 1 | cellular organisms |
| Bdellovibrionales bacterium GWA2_49_15 | 1 | cellular organisms |
| Mariprofundus ferrooxydans PV−1 | 1 | cellular organisms |
| Amantichitinum ursilacus | 1 | cellular organisms |
| Alphaproteobacteria bacterium RIFCSPHIGHO2_02_FULL_46_13 | 1 | cellular organisms |
| Rhodanobacter sp. B04 | 1 | cellular organisms |
| Acholeplasma laidlawii PG−8A | 1 | cellular organisms |
| Solirubrobacterales bacterium 67−14 | 1 | cellular organisms |
| Collinsella sp. CAG:289 | 1 | cellular organisms |
| Rubrobacter xylanophilus DSM 9941 | 1 | cellular organisms |
| Ferrimicrobium acidiphilum DSM 19497 | 1 | cellular organisms |
| Streptomyces cattleya NRRL 8057 = DSM 46488 | 1 | cellular organisms |
| Ardenticatena maritima | 1 | cellular organisms |
| Caldilinea aerophila DSM 14535 = NBRC 104270 | 1 | cellular organisms |
| Ktedonobacter sp. 13_2_20CM_2_56_8 | 1 | cellular organisms |
| Dehalococcoidia bacterium SM23_28_2 | 1 | cellular organisms |
| Longilinea arvoryzae | 1 | cellular organisms |
| Sphaerobacter thermophilus DSM 20745 | 1 | cellular organisms |
| Roseiflexus sp. RS−1 | 1 | cellular organisms |
| Fimbriimonas ginsengisoli Gsoil 348 | 1 | cellular organisms |
| Chthonomonas calidirosea T49 | 1 | cellular organisms |
| Deinococcus peraridilitoris DSM 19664 | 1 | cellular organisms |
| Peptoniphilus asaccharolyticus DSM 20463 | 1 | cellular organisms |

| Species | PS | Phylostratum name |
|---|---|---|
| Limnochorda pilosa | 1 | cellular organisms |
| Dialister micraerophilus DSM 19965 | 1 | cellular organisms |
| Eubacterium sp. CAG:252 | 1 | cellular organisms |
| Streptococcus dysgalactiae subsp. equisimilis RE378 | 1 | cellular organisms |
| Gemmatimonadaceae bacterium 4484_173 | 1 | cellular organisms |
| Chitinispirillum alkaliphilum | 1 | cellular organisms |
| Chitinivibrio alkaliphilus ACht1 | 1 | cellular organisms |
| Fibrobacter sp. UWH6 | 1 | cellular organisms |
| Lentisphaera araneosa HTCC2155 | 1 | cellular organisms |
| Criblamydia sequanensis CRIB−18 | 1 | cellular organisms |
| Phycisphaera mikurensis NBRC 102666 | 1 | cellular organisms |
| Paludisphaera borealis | 1 | cellular organisms |
| Kiritimatiella glycovorans | 1 | cellular organisms |
| Methylacidiphilum infernorum V4 | 1 | cellular organisms |
| Coraliomargarita sp. CAG:312 | 1 | cellular organisms |
| Pedosphaera parvula Ellin514 | 1 | cellular organisms |
| Chthoniobacter flavus Ellin428 | 1 | cellular organisms |
| Campylobacter concisus | 1 | cellular organisms |
| Archangium gephyra | 1 | cellular organisms |
| Ignavibacteria bacterium RBG_16_36_9 | 1 | cellular organisms |
| Chlorobium ferrooxidans DSM 13031 | 1 | cellular organisms |
| Phaeodactylibacter xiamenensis | 1 | cellular organisms |
| Chitinophaga eiseniae | 1 | cellular organisms |
| Cytophagaceae bacterium SCN 52−12 | 1 | cellular organisms |
| Porphyromonas gingivalis W83 | 1 | cellular organisms |
| Sphingobacterium spiritivorum ATCC 33861 | 1 | cellular organisms |
| Flavobacterium sp. A45 | 1 | cellular organisms |
| Theionarchaea archaeon DG−70 | 1 | cellular organisms |
| Hadesarchaea archaeon DG−33−1 | 1 | cellular organisms |
| Methanocella arvoryzae MRE50 | 1 | cellular organisms |
| Methanopyrus kandleri AV19 | 1 | cellular organisms |
| Archaeoglobus sulfaticallidus PM70−1 | 1 | cellular organisms |
| Pyrococcus furiosus DSM 3638 | 1 | cellular organisms |
| methanogenic archaeon ISO4−H5 | 1 | cellular organisms |
| Natrinema pellirubrum DSM 15624 | 1 | cellular organisms |
| Methanococcus aeolicus Nankai−3 | 1 | cellular organisms |
| Methanothermobacter thermautotrophicus str. Delta H | 1 | cellular organisms |
| Candidatus Methanohalarchaeum thermophilum | 1 | cellular organisms |
| Candidatus Nitrososphaera gargensis Ga9.2 | 1 | cellular organisms |
| Thermofilum adornatus | 1 | cellular organisms |
| Aphanomyces astaci | 2 | Eukaryota |
| Oryza sativa Indica Group | 2 | Eukaryota |

| Species | PS | Phylostratum name |
|---|---|---|
| Leishmania donovani BPK282A1 | 2 | Eukaryota |
| Acanthisitta chloris | 3 | Opisthokonta |
| Gorilla gorilla gorilla | 3 | Opisthokonta |
| Poecilia formosa | 3 | Opisthokonta |
| Anopheles arabiensis | 3 | Opisthokonta |
| Echinococcus granulosus | 3 | Opisthokonta |
| Mucor circinelloides f. circinelloides 1006PhL | 4 | Fungi |
| Nematocida parisii ERTm1 | 4 | Fungi |
| Encephalitozoon cuniculi GB−M1 | 4 | Fungi |
| Smittium culicis | 4 | Fungi |
| Agaricus bisporus var. burnettii JB137−S8 | 5 | Dikarya |
| Trametes cinnabarina | 5 | Dikarya |
| Cryptococcus gattii CA1280 | 5 | Dikarya |
| Moesziomyces antarcticus | 5 | Dikarya |
| Microbotryum intermedium | 5 | Dikarya |
| Neolecta irregularis DAH−3 | 6 | Ascomycota |
| Pneumocystis jirovecii RU7 | 6 | Ascomycota |
| Protomyces lactucaedebilis | 6 | Ascomycota |
| Saitoella complicata NRRL Y−17804 | 6 | Ascomycota |
| Schizosaccharomyces cryophilus OY26 | 6 | Ascomycota |
| Bipolaris maydis ATCC 48331 | 7 | Saccharomyceta |
| Zymoseptoria tritici ST99CH_1A5 | 7 | Saccharomyceta |
| Acremonium chrysogenum ATCC 11550 | 7 | Saccharomyceta |
| Neurospora tetrasperma FGSC 2508 | 7 | Saccharomyceta |
| Blastomyces dermatitidis ATCC 18188 | 7 | Saccharomyceta |
| Candida albicans SC5314 | 8 | Saccharomycetales |
| Candida arabinofermentans NRRL YB−2248 | 8 | Saccharomycetales |
| [Candida] auris | 8 | Saccharomycetales |
| Cyberlindnera fabianii | 8 | Saccharomycetales |
| Hanseniaspora guilliermondii | 8 | Saccharomycetales |
| Saccharomycetaceae sp. 'Ashbya aceri' | 9 | Saccharomycetaceae |
| Eremothecium gossypii ATCC 10895 | 9 | Saccharomycetaceae |
| Kazachstania africana CBS 2517 | 9 | Saccharomycetaceae |
| Kluyveromyces dobzhanskii CBS 2104 | 9 | Saccharomycetaceae |
| Lachancea dasiensis CBS 10888 | 9 | Saccharomycetaceae |
| Saccharomyces eubayanus | 10 | Saccharomyces |
| Saccharomyces uvarum | 10 | Saccharomyces |
| Saccharomyces arboricola | 11 | s5 |
| Saccharomyces kudriavzevii | 12 | s4 |
| Saccharomyces mikatae | 13 | s3 |
| Saccharomyces paradoxus | 14 | s2 |
| Saccharomyces cerevisiae | 15 | orphan |
